# Keratins coordinate tissue spreading by balancing spreading forces with tissue material properties

**DOI:** 10.1101/2025.02.14.638262

**Authors:** Suyash Naik, Yann-Edwin Keta, Kornelija Pranjic-Ferscha, Édouard Hannezo, Silke Henkes, Carl-Philipp Heisenberg

## Abstract

For tissues to spread, they must be deformable while maintaining their structural integrity. How these opposing requirements are balanced within spreading tissues is not yet well understood. Here, we show that keratin intermediate filaments function in epithelial spreading by adapting tissue mechanical resilience to the stresses arising in the tissue during the spreading process. By analysing the expansion of the enveloping cell layer (EVL) over the large yolk cell in early zebrafish embryos *in vivo*, we found that keratin network maturation in EVL cells is promoted by stresses building up within the spreading tissue. Through genetic interference and tissue rheology experiments, complemented by a vertex model with mechanochemical feedback, we demonstrate that stress-induced keratin network maturation in the EVL increases tissue viscosity, which is essential for preventing tissue rupture. Interestingly, keratins are also required in the yolk cell for mechanosensitive actomyosin network contraction and flow, the force-generating processes pulling the EVL. These dual mechanosensitive functions of keratins enable a balance between pulling force production in the yolk cell and the mechanical resilience of the EVL against stresses generated by these pulling forces, thereby ensuring uniform and robust tissue spreading.

## Introduction

Epithelial cell layer spreading is a core feature of multiple developmental and disease-related processes. In *Drosophila* development, for instance, the spreading of the epidermis leads to the closure of an opening at the dorsal side of the embryo^1,2^. Likewise, in wound healing, the epidermal cell layer spreads, and fusion closes the wound^3,4^. Various cellular processes have been proposed to contribute to epithelial cell layer spreading, including cell spreading, cell migration, oriented cell division, and cell intercalation^5,6^. These processes can generate the mechanical forces driving active tissue expansion and determine the aptitude of tissues to undergo spreading.

Zebrafish embryo morphogenesis is initiated by the spreading of the blastoderm over the large yolk cell in a process named epiboly^7^. During epiboly, the EVL, a simple squamous epithelial cell layer, formed at the surface of the blastoderm, undergoes massive spreading to eventually engulf the entire yolk cell at the end of epiboly (Figure 1A)^8,9^. EVL spreading has been shown to depend on the formation and contraction of a large actomyosin band positioned within a thin cytoplasmic layer at the surface of the yolk cell, the yolk syncytial layer (YSL)^10^. This actomyosin band within the YSL forms around the entire circumference of the yolk cell close to where the leading edge of the EVL contacts the YSL, and its contraction and flow are thought to generate the mechanical forces pulling the EVL over the yolk cell (Figure 1A)^10,11^. Both active spreading of EVL cells and oriented EVL cell divisions have been implicated in facilitating EVL spreading by modulating EVL surface tension^12,13^. However, EVL morphogenesis not only relies on the generation and transmission of active forces within the EVL and YSL, but also on changes in the material properties of the tissue resisting such forces^14^. Yet, the molecular and cellular mechanisms that determine EVL material properties and how they are spatiotemporally coupled to changes in active force production remain unclear. Understanding these mechanisms requires a closer examination of the cytoskeletal components within epithelial cells, particularly those that have been implicated in modulating tissue material properties.

**Figure 1:**
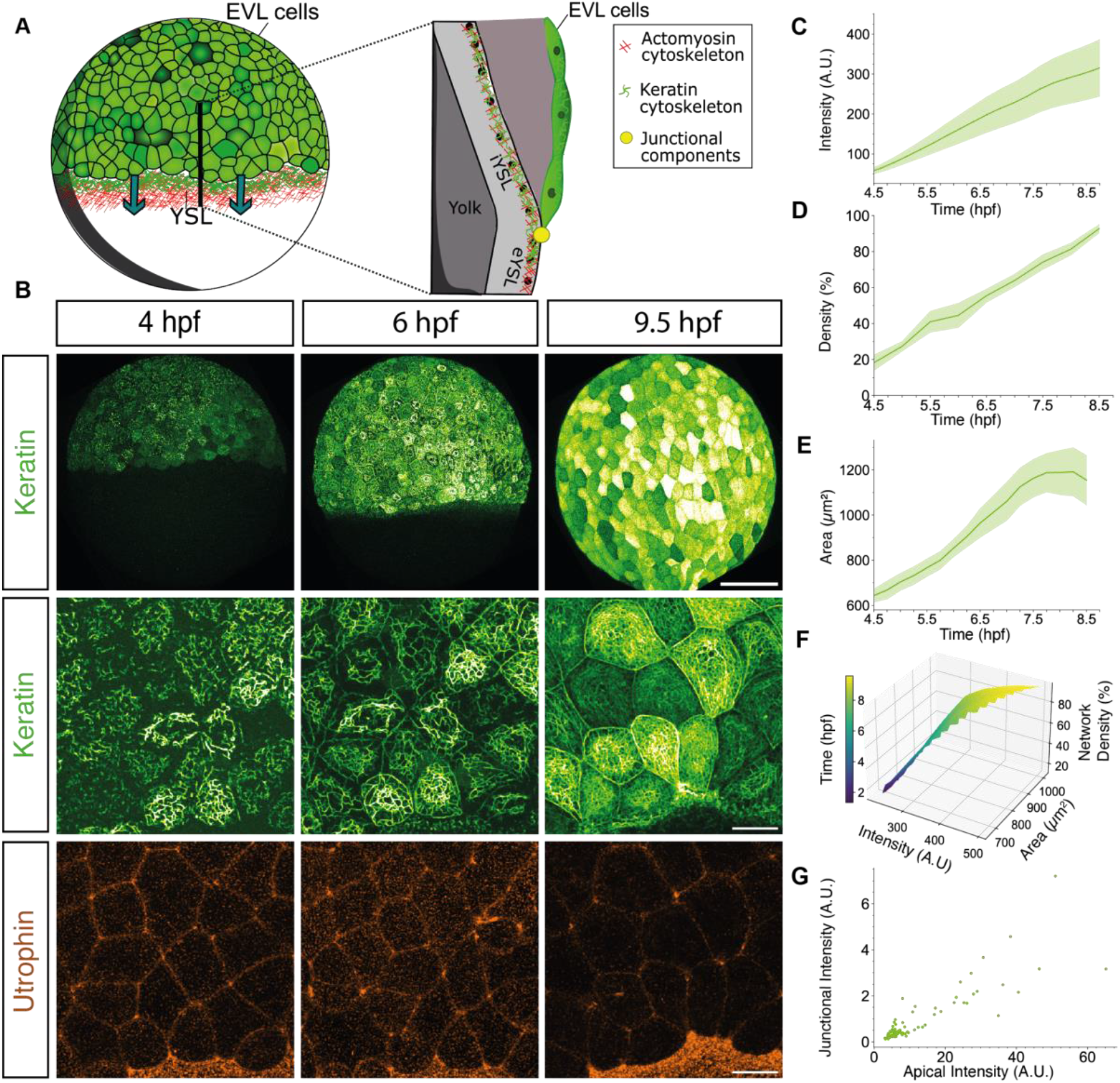
Keratin network maturation in the EVL. (A) Left: schematic representation of a zebrafish embryo at 60% epiboly stage (6 hpf) showing the expression and localization of keratin within the EVL and YSL (together with actin). Arrows mark the direction of EVL epiboly movements. Right: cross-section of the region outlined in the left panel marking the expression and localization of keratin, actin (within the YSL) and junctional components linking the margin of the EVL to the YSL. (B) Maximum intensity projection images of keratin (top and middle rows) and actin (Utrophin; bottom row)) expression in Tg*(krt18:Krt18-GFP)* (keratin) and Tg*(acbt2:Utrophin-mcherry)* (actin) embryos showing the progression of keratin expression within the EVL and YSL during epiboly (4 - 9.5 hpf). (C) Averaged keratin intensity within the EVL in Tg*(krt18:Krt18-GFP)* embryos as a function of time during epiboly (4 - 9.5 hpf). N=3 experiments; n=5 embryos. Error bars as ribbons SD of mean. (D) Averaged density of the keratin network in Tg*(acbt2:Utrophin-mcherry, krt18:Krt18- GFP)* embryos as a function of time during epiboly (4 - 9.5 hpf). N=3 experiments; n = x embryos. Error bars as ribbons SD of mean. (E) Average apical cell area of individual EVL cells in Tg*(acbt2:Utrophin-mcherry, krt18:Krt18-GFP)* embryos as a function of time during epiboly (4 hpf- 8.5hpf). N=3 experiments; n = 4 embryos. Error bars as ribbons SD of mean. (F) 3-dimensional (3D) plot of keratin intensity, network density, and EVL cell area as a function of time during epiboly. N=3 experiments; n = 4 embryos. Spread of the surface (width) represents data spread, indicating their variability (SD of intensity and area). (G) Correlation of junctional and apical keratin intensity measured in individual EVL cells in Tg*(acbt2:Utrophin-mcherry, krt18:Krt18-GFP)* embryos at 60% epiboly stage (6hpf) N=3 experiments, n = 3 embryos.

Keratin intermediate filaments are the most abundant and diverse cytoskeletal components in epithelial cells^15,16^. They form bundled filaments from keratin type I and type II heterodimers laterally associating in an antiparallel fashion into apolar tetramers that again align and anneal longitudinally into unit-length filaments^17,18^. Keratin filaments are semi-flexible, stable and highly elastic, different from the rather stiff and rigid actin and microtubule filaments^19,20^. They can self-assemble into intricate subcellular networks, the precise organisation of which depends on their functional adaptation in different cell types^21^. Generally, the intracellular keratin network can organise into a rim network, supporting plasma membrane integrity and connecting desmosomal contacts, and a spoke network, surrounding the nucleus and transferring information from the cell exterior to the nucleus^22–24^. In polarised simple epithelia, such as the EVL, keratin networks are typically positioned near the apex where they are thought to function in resisting mechanical and chemical stresses and maintaining epithelial apicobasal polarity^25–27^. Keratins have been shown to be sensitive to mechanical forces by reorganizing and changing their mechanical properties upon stress application^19,28,29^. Yet, how keratin network mechanosensitivity functions in epithelial tissue morphogenesis remains unsettled^30,31^.

The role of epithelial keratins in development and disease is only beginning to be understood^32,33^. Keratin mutations can cause diseases that lower the resilience of epidermal tissues to mechanical stress^34^. Moreover, studies in mouse embryos have shown that keratins are required for trophectoderm specification and extra-embryonic tissue growth and expansion^30,31^. Here we show that keratin intermediate filaments are required for EVL spreading during zebrafish epiboly. They function in this process by balancing EVL tissue viscosity with the external pulling forces mediating EVL spreading. This balancing function of keratins enables the EVL to undergo uniform spreading without rupturing.

## Results

### Keratins are specifically expressed within the EVL and YSL during gastrulation

To explore the function of keratins during early zebrafish embryogenesis, we analysed the expression of different keratins in embryos from early blastula to late gastrula stages (4.5-8.5 hpf). Previous studies have shown that 13 type I keratins and 3 type II keratins are specifically expressed within the developing EVL during this period^35,36^. To identify the temporal keratin expression profiles within the EVL during epiboly, we used RT-qPCR to map the expression of three keratin type II (*keratin 4, 5, 8*) and one keratin type I (keratin 18), abundantly expressed within the EVL^36,37^. We found all of these keratins to be expressed already at early blastula stages, with their expression continuously increasing until the end of gastrulation (Figure Sup 1A). The expression was restricted to the epithelial cells as evidenced by fluorescence *in situ* hybridization for *keratin 8* mRNA (Figure Sup 1B). To determine the spatial distribution of keratins within the epibolizing embryo, we took advantage of *Tgkrt18: krt18-GFP* embryos expressing GFP-tagged keratin 18 under its endogenous promotor. Keratin 18 expression was first detected in EVL progenitor cells within the early gastrula (4.0 hpf) arranged in short, bundled and unconnected filaments located predominantly at the apical surface of these cells (Figure 1B, Video 1 and Figure Sup 1C). Keratin 18 continued to be selectively expressed within EVL cells until mid-gastrulation (5.5 hpf), when some additional expression was also detected within the forming YSL directly adjacent to the place where the leading edge of the EVL contacts the YSL (Figure 1B). During this period and continuing until the end of gastrulation, the apical network of keratin 18 filaments within EVL cells became increasingly dense and interconnected (Figure 1B), as evidenced by a continuous increase in keratin 18- GFP intensity and network density between 4.5 and 8.5 hpf (Figure 1C and D, and Video 1). This increase in keratin network intensity and density was accompanied by EVL cells increasing the apical area during epiboly (Figure 1E), suggesting a close temporal correlation between keratin network maturation and EVL cell spreading (Figure 1F). Initially, keratin filaments were predominantly localized to the apical surface of the EVL cells (Figure 1B and Sup 1C), but from 6 hpf onwards, additional keratin 18 expression was detected in a bundle-like configuration along the apical junction of EVL cells (Figure 1B and G), consistent with previous observation of keratins showing both junctional (‘rim’) and apical (‘spoke’) localization^22,30,31^.

Finally, to check whether keratins in EVL cells can organise into networks different from the keratin 18 network, we ubiquitous expressed a mCherry-tagged form of keratin 4 by mRNA injection at the 1-cell-stage. Interestingly, keratin 4-mCherry, previously shown to form different keratin dimers than keratin 18^21^, was selectively expressed within EVL cells and its subcellular expression pattern entirely co-localized with the keratin 18 network (Figure Sup 1D), suggesting that different keratin dimers are part of the same network within EVL cells. Of note, EVL cells typically showed different levels of keratin expression both on the mRNA and protein level, with cells initially expressing higher levels also showing an earlier network maturation (Figure 1B, 1G and Sup 1B). This eventually resulted in the EVL at later stages of gastrulation being composed of cells displaying keratin networks at clearly different stages of maturation (Figure 1B and 1G).

Collectively, these findings suggest that keratins are predominantly expressed within the EVL and adjacent YSL, forming an increasingly dense apical and junctional network from early blastula to late gastrula stages accompanying EVL cell spreading.

### Keratin network maturation is mechanosensitive

EVL epiboly movements are driven by a large actomyosin cable forming within the YSL and pulling the margin of the EVL over the yolk cell^10^. Given that keratin networks can reorganise under stress^29,38^, we speculated that the observed maturation of the keratin network within EVL cells might be facilitated by EVL tissue tension building up during the course of epiboly^10,39^. To test this possibility, we analysed whether and how EVL network maturation is affected in embryos where EVL tension is either increased or decreased. To modulate EVL tension, we expressed a constitutive active form of RhoA (CARhoA) specifically within the YSL, promoting YSL actomyosin contractility and pulling, or a constitutive active form of Myosin phosphatase (CAMypt) leading to reduced YSL contractility and pulling^11^. In CARhoA- expressing embryos, keratin 18 expression intensity and network density in EVL cells prematurely increased during the course of epiboly, while in CAMypt-expressing embryos, keratin expression and network maturation were delayed (Figure 2A-D, Video 2-5 and Figure Sup 2B). This keratin mechanosensitivity was detectable both when network maturation was analysed as a function of developmental time or degree of epiboly progression (Figure 2A-D and Figure sup 2B-C), suggesting that the effect of EVL tension on keratin network maturation is not just a secondary consequence of changes in EVL epiboly movements (Figure Sup 2D). To further test this possibility, we dissociated EVL cells at the beginning (4hpf) and the middle of epiboly (6.5hpf) to obtain individual cells (Figure Sup 2E). In these cells, keratin network maturation was arrested at the time of their dissociation, with individual cells from embryos at 4 hpf showing a sparse keratin network and cells dissociated at 6.5 hpf showing a dense keratin network localized around the nucleus (Figure Sup 2E).

**Figure 2:**
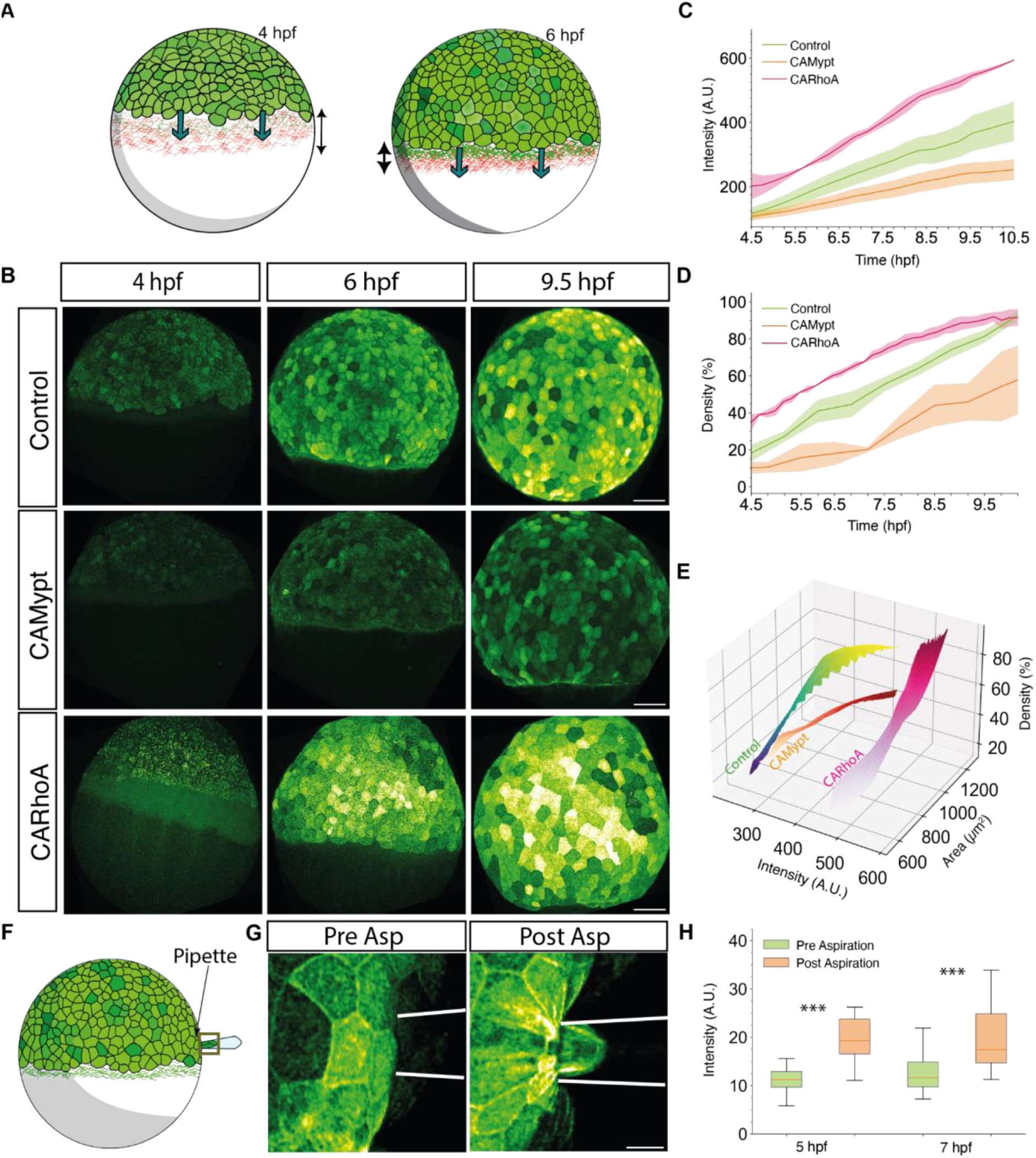
Pulling forces promote keratin expression within the EVL. (A) Schematic showing actomyosin contraction and flows within the YSL providing the mechanical forces pulling the EVL over the yolk cell during epiboly (for details see^10^) in a zebrafish embryo at 4hpf (left) and 6hpf (right). Green arrows, EVL epiboly movements; black arrows, actomyosin contraction within the YSL. (B) Maximum intensity projection images of keratin expression in representative Tg*(acbt2:Utrophin-mcherry, krt18:Krt18-GFP)* control embryos (YSL injection of 0.2% phenol red, top row) and embryos injected with 100pg *CAMypt* (middle row) and 50 pg *CARhoA* (bottom row) into either the YSL at 3.3 hpf (*CAMypt*) or marginal cells at 3.3 hpf (*CARhoA*) at sphere stage (4hpf, left column), shield stage (6 hpf, right column), and 90% epiboly stage (9.5hpf, right column). Scale bar: 100 µm (C) Plot of average keratin intensity as a function of time (hpf) in Tg*(krt18:KrtGFP)* control (green, N=3, n= 5 embryos), *CAMypt* (orange, n=3, n=5 embryos) and *CARhoA* mRNA injected embryos (pink, N=3, n=4 embryos) as described in (B). Error bars as ribbons SD of mean. (D) Plot of average density of keratin network as a function of time (hpf) in Tg*(actb2:Utrophin- mcherry, krt18:Krt18-GFP)* control (green, N=3, n= 6 embryos), *CAMypt* (orange, n=3, n=4 embryos) and *CARhoA* mRNA injected embryos (pink, N=3, n=3 embryos) as described in (B). Error bars as ribbon SD of mean of individual cells per replicate. (E) 3-dimensional (3D) plot of keratin intensity, network density and EVL cell area as a function of time (hpf) in Tg*(actb2:Utrophin-mcherry, krt18:Krt18-GFP)* control (viridis, N=3, n= 3 embryos), *caMypt* (orange, n=3, n=3 embryos) and *caRhoA* mRNA injected embryos (WPk, N=3, n=3 embryos) as described in (B). Spread of the surface (width) represents data spread, indicating their variability (SD of intensity and area). (F) Schematic showing pipette aspirations of the EVL of a 70% epiboly (7hpf) stage embryo where the regions of interest within the pipette and outside of it are marked as yellow boxes. (G) Maximum intensity projection images of keratin localization and intensity within the EVL before (left) and after (right) aspiration with a pipette in a representative Tg*(krt18:KrtGFP)* embryo. White lines outline the boundary of the pipette. Scale bar: 10 µm. (H) Box plot of keratin intensity within the EVL before (green) and after (orange) aspiration either within the pipette (right box in schematic) or outside of it (left box in schematic) in Tg*(actb2:Utrophinmcherry,krt18:Krt18GFP)* embryos at 5 and 7hpf. Boxes represent quartiles for the data, dots outliers (pvalues: ***<0.001 Wilocoxon test).

Finally, to directly assess the effect of EVL tension on keratin network maturation, we locally increased EVL tension in 5 and 7 hpf embryos by aspirating the EVL using a micropipette and analysing resultant changes in EVL network maturation. We found that keratin filaments showed increased accumulation in EVL cells upon aspiration (Figure 2F-H, Video 8), further supporting the notion that keratin network maturation within the EVL cells is promoted by EVL tension building up during the course of epiboly.

### Keratins regulate EVL tissue properties

Keratins have previously been implicated in resisting mechanical stresses in epithelial cells^25^. Thus, the mechanosensitive coupling between EVL keratin network maturation and pulling force generation within the YSL might constitute a mechanism to protect the EVL tissue against excessive deformations by mechanically strengthening it proportionally to the force exerted on it.

To test this possibility, we knocked down/out the expression of the two keratin type II genes (*keratin 4 and 8*) primarily expressed within EVL cells^36^, reasoning that in the absence of keratin type II expression, no keratin dimers can be formed within the EVL and thus keratin network formation should be defective. Ubiquitous knock-down/out of *keratin 4* and *8* expression by using *morpholinos* and Crispr-Cas9 or interference with keratin network formation by overexpressing a dominant negative version of Keratin 18 (DNkeratin18)^40^, led to strongly diminished keratin 18 expression and network formation within EVL cells (Figure 3A-C, Video 6-7 and Sup 3A-B). Loss of the keratin network in all these perturbations lead to delayed EVL epiboly movements (Figure 3D, Video 6 and 7 and Figure Sup 3C) and frequent rupturing of the EVL towards the end of gastrulation, causing embryo lethality (Figure Sup 3D-E).

**Figure 3:**
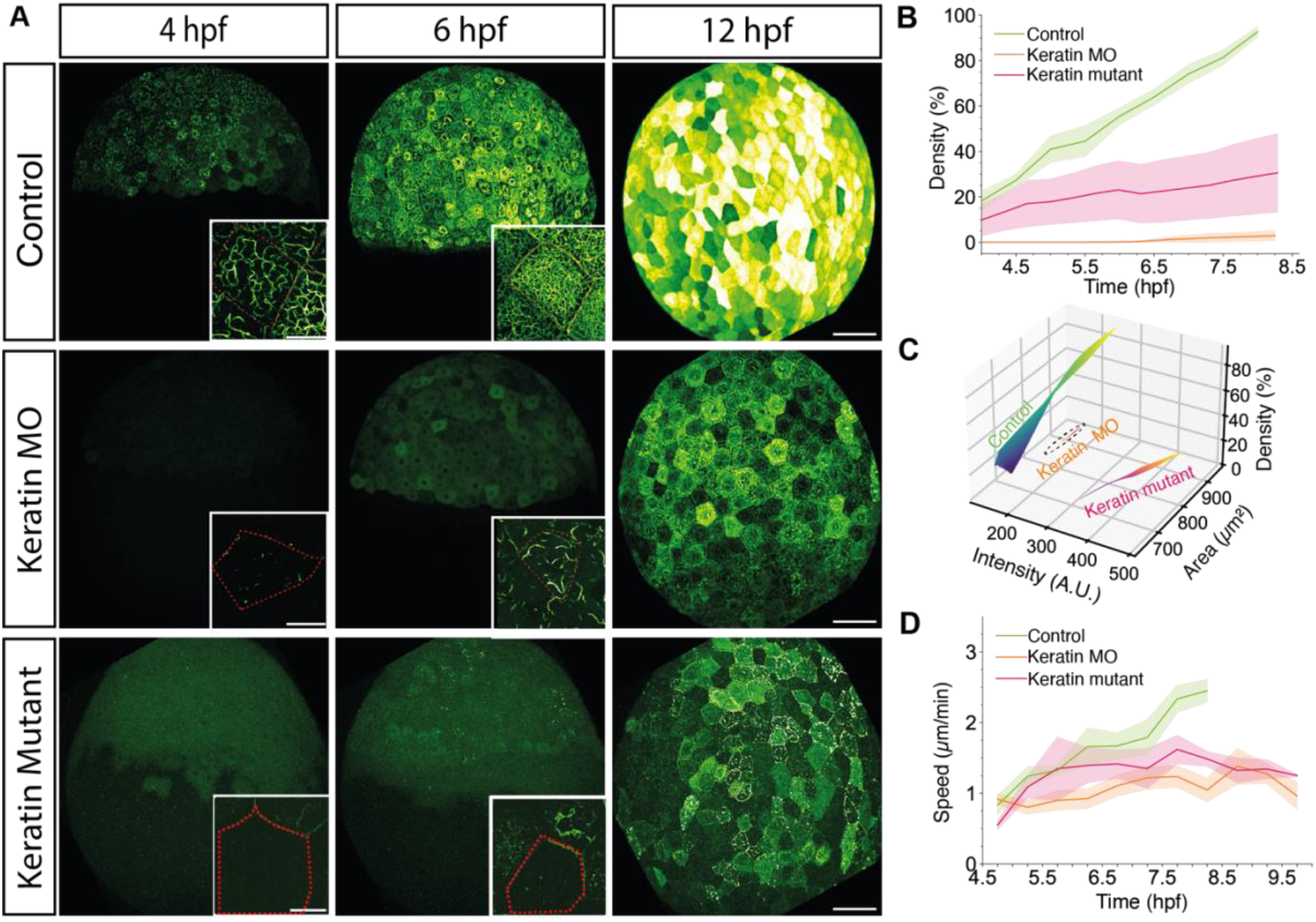
Loss of keratin expression diminishes EVL epiboly movements. (A) Maximum intensity projection images of keratin expression in representative Tg*(krt18: Krt18GFP)* embryos at sphere stage (4hpf, left column), shield stage(6phf, middle column) and bud stage (10hpf, right column) with insets on the right lower corner showing single cells with their boundary marked by a red line injected at the one-cell stage with 2ng control MO (top row), 1ng *krt4* plus 1ng *krt8* MO (middle row), or with TraCr *krt4* and *krt8* gRNA (*krt4/8* crispant F0; bottom row). Scale bar: 100 µm (B) Plot of averaged density of keratin network in individual EVL cells as a function of time (hpf) during epiboly in Tg*(actb2:Utrophinmcherry, krt18:Krt18GFP)* embryos injected at the one-cell stage with 2ng control MO (top row; green, N=4, n=4 embryos), 1ng *krt4* plus 1ng *krt8* MO (middle row; orange, N=4, n=4 embryos), or with TraCr *krt4* and *krt8* gRNA (*krt4/8* crispant F0; bottom row; pink, N=2, n=6 embryos). Error bars as ribbon SD of the mean of individual cells per replicate. (C) 3-dimensional (3D) plot of keratin intensity, network density and area of EVL cells as a function of time (hpf) during epiboly in Tg*(actb2:Utrophinmcherry,krt18:Krt18GFP)* embryos injected at the one-cell stage with 2ng control MO (top row; green, N=4, n=4 embryos), 1ng *krt4* plus 1ng *krt8* MO (middle row; orange, N=4, n=4 embryos), or with TraCr *krt4* and *krt8* gRNA (*krt4/8* crispant F0; bottom row; pink, N=2, n=6 embryos). Spread of the surface (width) represents data spread, indicating their variability (SD of intensity and area). (D) Plot of EVL epiboly movement speed as a function of time (hpf) during epiboly starting at sphere stage (4hpf) until late epiboly stages (9 hpf) in Tg*(actb2: Utrophin-mcherry, krt18:Krt18GFP)* embryos injected at the one-cell stage with 2ng control MO (top row; green, N=3, n=4 embryos), 1ng *krt4* plus 1ng *krt8* MO (middle row; orange, N=3, n=4 embryos), or with TraCr *krt4* and *krt8* gRNA (*krt4/8* crispant F0; bottom row; yellow, N=2, n=4 embryos). Error bars as SD of the mean of replicates.

To understand how keratins function in EVL epiboly movements, we analysed changes in EVL cell shapes and rearrangement in response to the pulling forces from the YSL during epiboly. Comparing wild-type with *keratin 4/8* morphant embryos revealed that EVL cells in wild-type embryos coordinately elongated along the animal-vegetal (AV) axis of the gastrula, the axis of EVL spreading, while no such coordinated cell elongation was detectable in morphant embryos (Figure 4A-B). This points at the possibility that tissue material properties, determining force transduction and mechanical resilience, might be altered by keratin network formation.

**Figure 4:**
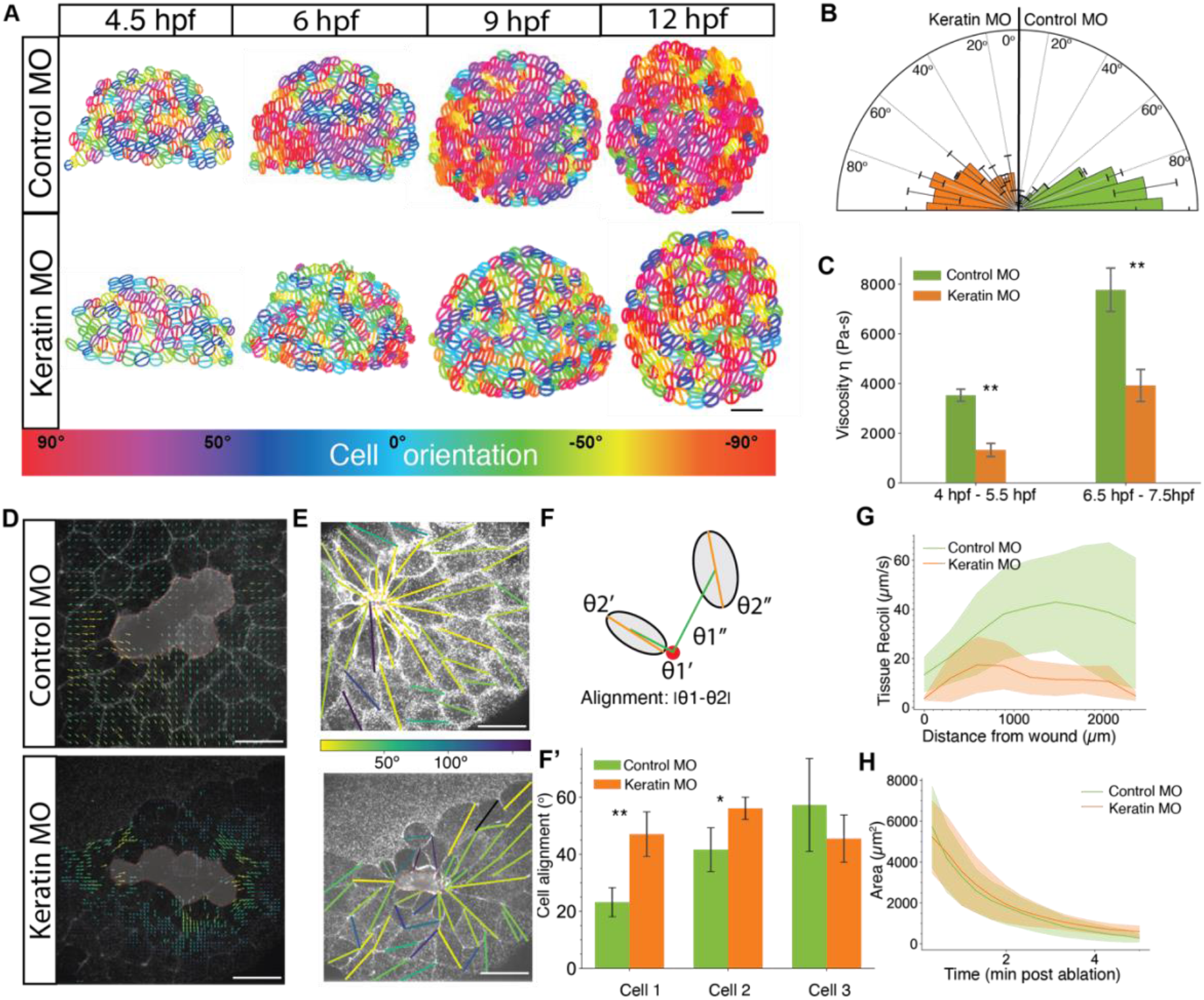
Mechanical force percolation within the EVL is dependent on keratin expression. (A) Plots of EVL cell orientations with ellipses representing shape descriptors (long and short axis) of individual EVL cells with the line in the middle marking the orientation of the long axis at consecutive stages during epiboly (4.5, 6, 9, 12 hpf) in curvature corrected Tg*(actb2:Utrophin-mcherry, krt18:Krt18GFP)* embryos injected at the one-cell stage with 2 ng control MO (top row) or 1ng *krt4* plus 1ng *krt8* MO (middle row). Each cell is colour-coded according to the orientation of the axis (hsv) as shown in the colour bar at the bottom (Red: AV axis orientation, blue: dorsoventral/DV orientation). (B) Rose plot of EVL cell orientations in curvature corrected Tg*(actb2:Utrophinmcherry,krt18:Krt18GFP)* embryos at 6 hpf injected at the one-cell stage with 2 ng control MO (B; N=4, n=93 cells) or 1ng *krt4* plus 1ng *krt8* MO (C; N=4 n= 84 cells). (C) Bar plots of tissue viscosity measured at the EVL margin using micropipette aspiration at 4-6 hpf and 6-7.5 hpf in Tg*(actb2: Utrophin-mcherry, krt18:Krt18GFP)* injected at the one- cell stage with 2 ng control MO (B; N=5, n=37 embryos) or 1ng *krt4* plus 1ng *krt8* MO (C; N=5, n=37 embryos). (D) Representative quiver plots of EVL tissue recoil flow velocities after cell ablation/wounding (wound centre marked by a red dot) in Tg*(actb2:Utrophinmcherry)* embryos at 6 hpf injected with 2 ng control MO (top) or 1ng *krt4* plus 1ng *krt8* MO (bottom). The arrows show the local velocity coloured according to the magnitude (viridis). (E) Average radial recoil velocity of the tissue flow plotted as a function of distance from would centre (0 µm) in Tg*(actb2:Utrophinmcherry)* embryos injected with 2 ng control MO (top) or 1ng *krt4* plus 1ng *krt8* MO (bottom) at successive time points after cell ablation (from yellow to blue; N=23 embryos). (F) Left panels: EVL cells in Tg*(actb2:Utrophinmcherry)* embryos injected with 2 ng control MO (top) or 1ng *krt4* plus 1ng *krt8* MO (bottom) and overlaid with lines representing their orientation (line angle) and alignment (colour-coded according to the alignment angle; reversed viridis). Right upper panel: schematic showing how cell alignment was determined by measuring the angle from the wound centre (ϴ1, green lines) and the cell longest cell axis (ϴ2, orange lines). The alignment was calculated by determining the magnitude of the difference between the angle from the wound centre and the cell longest cell axis (|ϴ1-ϴ2|) in successive rows of cells around the wound centre. Right lower panel: bar plot (bottom) of EVL cell alignment in Tg*(actb2:Utrophinmcherry)* embryos injected with 2 ng control MO (green) or 1ng *krt4* plus 1ng *krt8* MO (orange) upon control MO injection (green) in successive cell rows (cell 1 - cell 3) around the wound center. (N=23 embryos). (H) Plot of wound area as a function of time upon ablation of EVL cells in Tg*(actb2:Utrophinmcherry)* embryos injected with 2 ng control MO (green) or 1ng *krt4* plus 1ng *krt8* MO (orange). N=23 embryos.

To investigate this possibility, we measured the material properties of EVL tissue at early and mid-gastrulation (5 and 7 hpf) using micropipette aspiration. Analysis of the flow profile of the EVL into the pipette revealed a linear response, consistent with viscous properties, which allowed us to assess the viscosity of EVL tissue (Figure Sup 3F). Comparing the viscosity of EVL tissue between early and mid-gastrulation stages showed a significant increase in viscosity at mid-gastrulation, coinciding with the maturation of the keratin network (Figure 4C). This suggests that keratin network maturation during epiboly contributes to the increase in EVL tissue viscosity. To test this function of keratins more directly, we performed EVL micropipette aspiration experiments in wild-type and *keratin 4/8* morphant embryos. At early gastrulation, when differences in keratin network formation between wild-type and morphant embryos were still relatively small (Figure 1B), we observed a slight reduction in viscosity in *keratin 4/8* morphant embryos compared to wild-type (Figure 4C). In contrast, at mid- gastrulation (Figure 4C, Video 8), when the differences in keratin network organization between morphant and wild-type embryos became more pronounced (Figure 1B), viscosity was significantly reduced in the morphant embryos. This supports the idea that keratin network maturation enhances EVL tissue viscosity.

To test whether this effect is due to keratin expression specifically within EVL cells, we knocked down *keratin 4/8* expression within the YSL by injection of *keratin 4/8* morpholinos directly into the forming YSL at the 512-cell stage. Interestingly, this did not cause any detectable changes in coordinated EVL cell elongation along the AV axis, suggesting that keratins control EVL cell shape changes in a tissue-autonomous manner (Figure Sup 4A-C). To further challenge this conclusion, we developed an EVL wound healing assay, allowing us to monitor autonomous EVL spreading during wound closure^41,42^. For wounding the EVL, we ablated 3-5 neighbouring cells at random positions within the EVL in mid-gastrulation stage embryos (7 hpf) and observed how the coordinated spreading of the neighbouring cells extruded the ablated cells. In control embryos upon ablation, a supracellular actin cable formed at the leading edge of EVL cells neighbouring the ablated cells (Figure 4D, Video 9-10). This was accompanied by the highly coordinated movement and spreading of EVL cells towards the site of cell ablation, eventually leading to the extrusion of the ablated cells and closure of the wounding site (Figure 4G). During this process, not only the leading edge EVL cells elongated towards the wounding site, but also cells further away from the ablation site, resulting in a highly coordinated long-range tissue spreading detected by tissue flow and cell alignment analysis (Figure 4D-G). In contrast, EVL oriented cell elongation and tissue spreading were largely restricted to EVL cells directly neighbouring the wounding site in *keratin 4/8* morphant embryos (Figure 4D-G). This supports the notion that keratin expression within the EVL promotes tissue viscosity, leading to an increased length (hydrodynamic length) by which the tissue deforms in response to forces pulling at its margin.

### Keratins are required for mechanosensitive actomyosin contraction within the YSL

Given that keratin expression within the EVL is mechanosensitive, we hypothesised that this behaviour might constitute a feedback mechanism balancing EVL tissue viscosity with the forces pulling on its margin, thereby setting the rate of EVL epiboly movements. To address this hypothesis, we first turned to our EVL wound closure assay, allowing us to analyze the tissue-autonomous role of keratins in EVL spreading. Comparing the rate of wound closure between wild-type and *keratin 4/8* loss-of-function embryos revealed no apparent differences (Figure 4H), suggesting that - for the short timeframe of wound closure - reduced tissue viscosity in *keratin 4/8* loss-of-function embryos has no significant consequences on the rate of EVL spreading. To conceptualise how this behaviour can be explained by keratin expression within the EVL, we developed a vertex model^43,44^ of the EVL, enabling us to relate EVL tissue spreading dynamics to mechanosensitive keratin expression. To this end, we wrote the energy 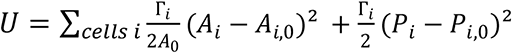 which penalises deviations of the areas *A_i_* and perimeters *P_i_* of the cells *i* from their target values *A*_*i*,_*_0_* and 𝑃_*i*,_*_0_*, respectively, with cell stiffness constants 𝛤_*i*_. In order to make the tissue viscoelastic, we relaxed the target area *A*_*i*,_*_0_* from their initial average value *A_0_* following 𝜏_*i*_ 𝑑*A*_*i*,_*_0_*/𝑑𝑡 = −(*A*_*i*,_*_0_* − *A*_*i*_) and scale 𝑃_*i*,_*_0_* to work at a constant shape parameter 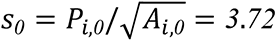, a value where the tissue is predicted to be still rigid^43^.

We then introduce keratin mechanosensitivity as follows (Figure 5A): as shown in Figure 1B, keratin forms a network within each cell that is initially floppy, but reaches a percolation threshold around 𝐾_𝑡*ℎ*_ = *150*. Above this, we assumed that it acts in parallel with the other mechanical components of the cell, increasing both cell stiffness and relaxation time proportionally, with 𝛤_*i*_ = Γ(*1* + 𝛽 𝛥𝐾_*i*_) and 𝜏_*i*_ = τ(*1* + 𝛽 𝛥𝐾_*i*_), where 𝛥𝐾_*i*_ = 𝑚𝑎𝑥(*0*, 𝐾_*i*_ − 𝐾_𝑡*ℎ*_). To include keratin mechanosensitivity, we further assumed that its assembly from the cytosol increases with pressure. To account for this, we used a simple, linear model of the evolution of the keratin concentration 𝜏_𝐾_ 𝑑𝐾_*i*_/𝑑𝑡 = 𝛼 𝑚𝑎𝑥(*0*, 𝑝_*i*_) − 𝐾_*i*_, where keratin dissociates with time scale 𝜏_𝐾_ (note that other nonlinear rheology models have also been used to model the effect of keratin^29^). Mechanosensitivity enters through the pressure 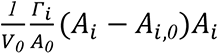, which takes into account the 3D cell volume 𝑉, with a coupling constant 𝛼 (see SI section 2). In steady-state, this corresponds to a linear keratin-pressure relation 𝐾_*i*_ = 𝛼 𝑝_*i*_ assuming that the pressure is positive (for a summary of this mechanochemical feedback loop see Figure 5B). Finally, to estimate the parameters for simulating EVL wound closure and epiboly, we used the EVL aspiration experiments described above (Figure 2F–H and 4C) together with a mean field version of the mechanical model (see SI section 1). In particular, we separately fitted the low-keratin initial response and the high-keratin release curve in the aspiration experiments for obtaining estimates of 𝜏_𝐾_, 𝛼, 𝛤, and 𝜏. We have chosen 𝛽 = *0*.*005* for the wild-type model tissue such that the increase of keratin levels leads to a 2- to 3-fold increase of Γ_*i*_ and 𝜏_*i*_. Please see table S1 for the full list of fitted parameters.

**Figure 5:**
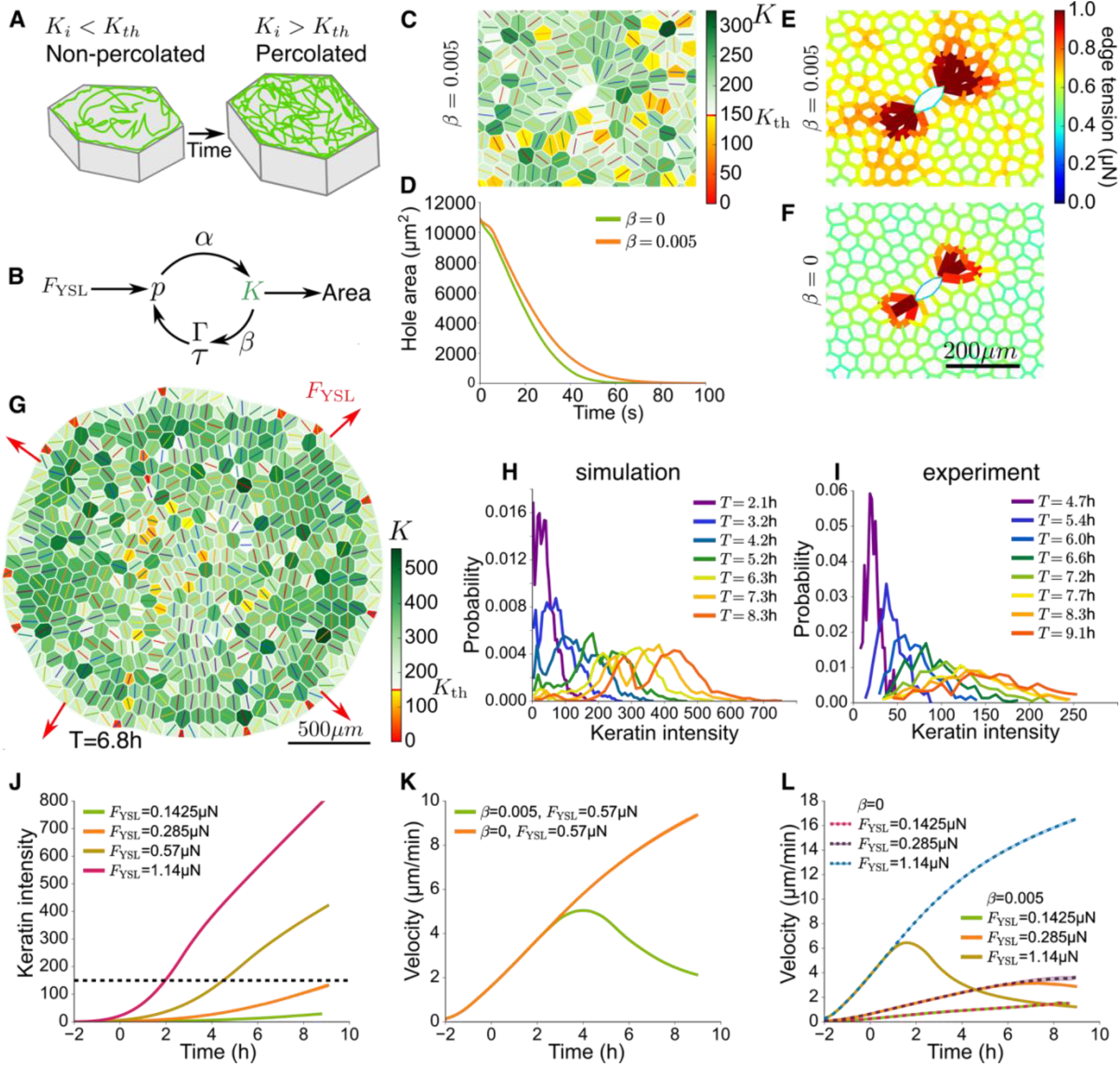
Vertex model of EVL spreading in the presence and absence of keratin expression. (A) Illustration of the keratin network in an non-percolated (mechanically inactive) and percolated (mechanically active) state. (B) Keratin mechanochemical feedback loop: pulling force 𝐹_𝑌𝑆𝐿_ causes pressure 𝑝 in the tissue that enhances keratin 𝐾 formation, which increases tissue stiffness 𝛤 and relaxation time 𝜏, which again increases pressure 𝑝. (C) Keratin expression at 𝑡 = *30*𝑠 in a EVL model tissue with periodic boundary conditions, where at 𝑡 = *0*𝑠 6 cells are removed and a tension of 3µN is applied on the edges surrounding the removal site (hole). (D) Area of the hole (shown in C) as a function of time after ablation in keratin deficient (𝛽 = *0*; green) and wild-type with (𝛽 = *0*.005; orange) EVL model tissues. (E,F) Edge tension network in a wild-type (E; 𝛽 = *0*.005) and keratin deficient (F; 𝛽 = *0*) EVL model tissue after ablation at 𝑡 = *30*𝑠. (G) Keratin expression in an EVL model tissue at mid-epiboly; lines are axes of cell orientation. (H) Distribution of keratin intensity within an EVL model tissue at consecutive timepoints during spreading (epiboly). (I) Distribution of keratin intensity within the EVL of a representative Tg*(krt18: Krt18GFP)* embryo at consecutive timepoints during epiboly (4.5 - 9.1 hpf). (J) Mean keratin expression for different values of 𝐹_𝑌𝑆𝐿_ ; orange corresponds to the wild-type value. (K) Tissue edge velocity for ‘wild-type’ 𝐹_𝑌𝑆𝐿_ (see also J) in keratin deficient (𝛽 = *0*; orange) and wild-type (𝛽 = *0*.005; green) model tissues. (L) Tissue edge velocity for 𝐹_𝑌𝑆𝐿_ values lower (brown, green) and higher (pink, orange) than the ‘wild-type’ value (see also J) in keratin deficient (𝛽 = *0*; brown, pink) and wild-type (𝛽 = *0*.005; green, orange) model tissues. Error bars for (J, K, L) have a width of 1 standard deviation computed over 10 independent runs.

Using this extended vertex model (see simulation library^45^), we first simulated the EVL wound closure behavior in wild-type and keratin deficient embryos. For this, we initialised a disordered tissue patch under tension, and then created a model wound by deleting several cells and adding a contractile cable to the wound perimeter (Figure 5C-F; for details see SI section 3). We observed that, similar to our experimental observations, tension was higher and more widely distributed around the wound in wild-type (𝛽 = *0*.*005*) compared to keratin deficient (𝛽 = *0*) tissues (Figure 5E-F), while the time required to close the wound remained largely unchanged between these conditions (Figure 5D). This close match between our experimental and theoretical observations supports the plausibility of keratin forming a mechanochemical feedback loop within the EVL.

To test whether we can also simulate the behaviour observed for EVL spreading during epiboly in wild-type and keratin-deficient embryos, we simulated epiboly by representing the EVL tissue as a disordered circular packing of N=529 cells, where the outer vertices are pulled outwards (Figure 5G). As the actual embryo is spherical, we defined the effective height of the model tissue 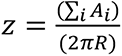, which is the height of its projection on a sphere of radius 𝑅=350µm, and we defined the tissue’s velocity 𝑣_𝑧_ as the derivative of this height with respect to time. To mimic the pulling of the EVL margin by actomyosin contraction and flow within the YSL, we applied an outward force 𝐹_𝑌𝑆𝐿_ on the edge of the simulated tissue with an amplitude which increases linearly with time, following the experimentally determined pulling force evolution within the YSL^10^. We then measured the mean keratin expression 𝐾 and edge velocity 𝑣_𝑧_ as a function of time and pulling forces (Figure 5J-L). By ramping the pulling force from *0* at the beginning to *0*.*57* 𝜇𝑁 (a value equivalent to the pulling force used in the aspiration experiments) at the end of the simulation, we obtained keratin expression values and tissue edge velocities matching our experimental observations (Figure 1C, 3D and 5K). Interestingly, we also found, similar to our experimental observations (Figure 5I), that keratin expression increased to heterogeneous levels in individual EVL cells in our simulations (Figure 5H), suggesting that this heterogeneous keratin expression is mechanically regulated.

To determine whether our model can also account for the experimentally observed changes in EVL dynamics when pulling forces within the YSL were either increased or decreased, we analysed the response of keratin expression and tissue edge velocity to variations in pulling forces (𝐹_𝑌𝑆𝐿_) in our simulations (Figure 5J, L). Consistent with experimental observations,

mean keratin expression increased when 𝐹_𝑌𝑆𝐿_ was upregulated and decreased when 𝐹_𝑌𝑆𝐿_ was downregulated (Figure 5J). Moreover, in line with the observed scaling of keratin expression, and thus tissue viscosity, with 𝐹_𝑌𝑆𝐿_, edge velocity in our simulations was not significantly affected by changes in 𝐹_𝑌𝑆𝐿_ (Figure 5L). However, these predictions did not align with our experimental findings at later stages of gastrulation, where both increasing and decreasing 𝐹_𝑌𝑆𝐿_ - by modulating actomyosin contractility within the YSL - slowed EVL epiboly movements (Figure supplement 2D). This suggests that changes in keratin expression, and consequently EVL tissue viscosity, cannot fully compensate for alterations in 𝐹_𝑌𝑆𝐿_ throughout gastrulation. A similar discrepancy between model predictions and experimental results emerged when comparing tissue edge velocity in wild-type and keratin-deficient embryos. While our simulations predicted a significant increase in edge velocity in keratin-deficient embryos compared to wild-type (Figure 5K) - as expected for a less viscous and more deformable EVL - experimental data showed a decrease in edge velocity in keratin-deficient embryos (Figure 3D).

Interestingly, in our simulations, the reduction in EVL tissue edge velocity observed in keratin-deficient embryos could only be explained by a decrease in the pulling force 𝐹_𝑌𝑆𝐿_ (Figure 5L). This led us to speculate that keratins may have additional functions within the YSL in regulating 𝐹_𝑌𝑆𝐿_ . Supporting this hypothesis, both our findings and previous studies indicate that keratins are also expressed within the YSL (Figure 6A) and play a role in actomyosin network organization and mechanosensation^46,47^. To test this possibility, we analysed dynamic actomyosin reorganization within the YSL in the presence and absence of keratins. In wild-type embryos, retrograde flows of actin and myosin within the YSL led to the formation of a contractile actomyosin band, previously shown to generate the pulling forces that drive EVL spreading^10^. However, in embryos where keratin 4/8 was specifically knocked down within the YSL - achieved by injecting *keratin 4/8* morpholinos directly into the forming YSL at the 512- cell stage - these actomyosin flows were severely diminished and less aligned (Figure 6A, D, F).

**Figure 6:**
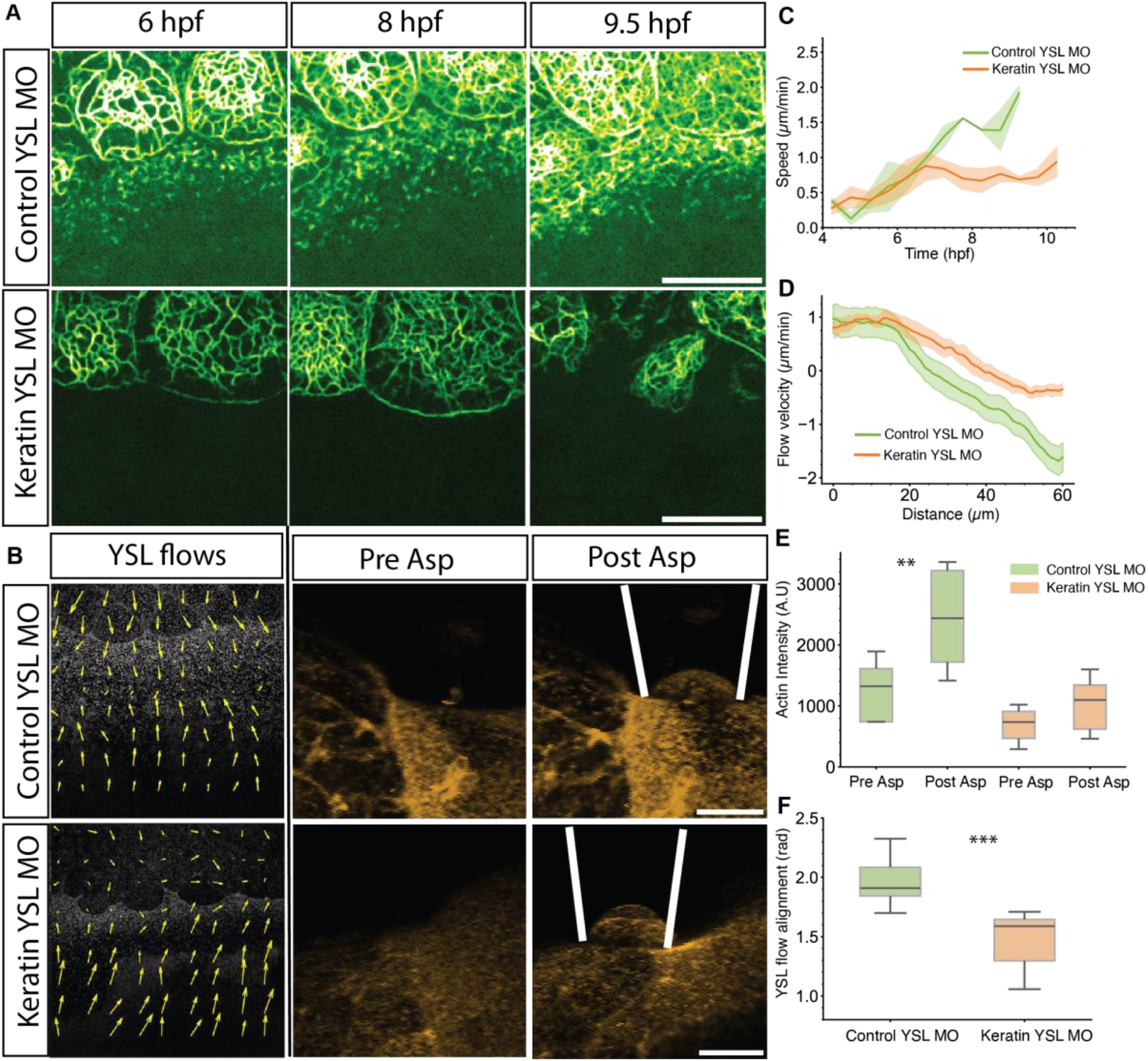
Actin flow alignment within the YSL is dependent on keratin expression. (A) Maximum intensity projections of the keratin network at the EVL-YSL boundary of Tg*(krt18:Krt18GFP)* embryos at shield (left column), 75% epiboly (middle column) and 95% epiboly (right column) injected with 2ng control MO (top row) or 1ng *krt4* plus 1ng *krt8* MO (bottom row) into the YSL at sphere stage (3.3 hpf). Scale bar, 25 µm (B) Maximum intensity projections of the actin network at the EVL-YSL margin of Tg*(actb2:Utrophinmcherry,krt18:KeratinGFP)* embryos at 6 hpf injected with 2ng control MO (top row) or 1ng *krt4* plus 1ng *krt8* MO (bottom row) into the YSL at sphere stage (3.3 hpf) before (Pre Asp, left column) and after (Post Asp, middle column) micropipette aspiration. Scale bar: 25 µm. Right column: Representative maximum intensity projections of actin and keratin in Tg*(actb2:Utrophinmcherry,krt18:Keratin18GFP)* at 3.3 hpf injected with 2ng control MO (top panel) or 1ng *krt4* plus 1ng *krt8* MO (bottom panel). Images are overlaid with quiver plots of retrograde actin flows within the YSL. Scale bar, 50 µm (C) Plot of EVL epiboly movement speed as a function of time (hpf) during epiboly in Tg*(actb2: Utrophin-mcherry, krt18:Krt18GFP)* embryos injected with 2ng control MO (green) or 1ng *krt4* plus 1ng *krt8* MO (orange) into the YSL at sphere stage (3.3 hpf). Error bars as ribbon SD of mean. N=4, n= 8 embryos. (D) Plot of retrograde actin flow velocity in the YSL as a function of distance from the EVL- YSL boundary in Tg*(actb2: Utrophin-mcherry, krt18:Krt18GFP)* embryos at shield stage (6 hpf) injected with 2ng control MO (green) or 1ng *krt4* plus 1ng *krt8* MO (orange) into the YSL at sphere stage (3.3 hpf). Error bars as ribbon SD of mean. N=3 n= 6 embryos. (E) Box plot of actin intensity in a fixed region of interest (ROI) within the YSL close to the point of micropipette aspiration before (pre Asp) and after (post Asp) aspiration in Tg*(actb2: Utrophin-mcherry, krt18:Krt18GFP)* embryos at 3.3 hpf injected with 2ng control MO (green boxes) or 1ng *krt4* plus 1ng *krt8* MO (orange boxes) into the YSL at sphere stage (3.3 hpf). N=4, n=13 embryos. Boxes represent quartiles of the data and error bars the spread (pvalues: ***<0.001,**<00.1 paired t test). (F) Box plot of actin flow alignment within the YSL at 6 hpf in Tg*(actb2: Utrophin-mcherry, krt18:Krt18GFP)* embryos injected with 2ng control MO (green boxes) or 1ng *krt4* plus 1ng *krt8* MO (orange boxes) into the YSL at sphere stage (3.3 hpf). Boxes represent quartiles of the data and error bars the spread. N=3, n=6 (***<0.001, paired t test).

To determine how keratins functionally interact with the actomyosin network within the YSL, we asked whether tension, generated at the EVL-YSL boundary and propagating into both the EVL and YSL, might not only promote keratin network formation within the EVL, but also increase actomyosin contraction and flows within the YSL. We further speculated that such potential effect of mechanical tension on actomyosin contraction and flow within the YSL would rely on keratin network formation within the YSL, analogous to previous reports linking keratins to actomyosin mechanosensation^46,47^. To test these possibilities, we performed micropipette aspiration experiments of the YSL and monitored resultant mechanosensitive changes in actomyosin network formation in wild-type and keratin loss-of-function embryos. While in wild-type embryos, YSL pulling led to a local increase in the intensity of actin next to the pipette (Figure 6B and E), in embryos where *keratin 4/8* was specifically knocked down within the YSL, no such pronounced upregulation was observed (Figure 6B, E). This suggests that keratins are required for the generation of pulling forces within the YSL driving EVL epiboly movements by facilitating actin mechanosensation within the YSL.

## Discussion

Our findings suggest that keratins function in EVL spreading during epiboly by coupling tissue contractility and tension to tissue viscosity and connectivity. During epiboly, actomyosin contraction and flow within the YSL generate the forces pulling the margin of the EVL towards the vegetal pole of the gastrula. These pulling forces not only induce stress within the EVL, but also within the YSL to which the EVL is mechanically coupled at its margin. In the EVL, buildup of stress triggers keratin network maturation, which again increases EVL tissue viscosity, thereby ensuring that the tissue remains intact when stress rises during epiboly. In the YSL, keratins are required for stress-dependent actin network accumulation and efficient contraction needed for pulling the EVL over the yolk cell. This dual mechanosensitive function of keratin, coupling EVL viscosity to YSL contractility, secures robust and efficient EVL spreading during epiboly.

Keratins have previously been shown to be stress-responsive and display important functions in mechanical tissue resilience and spreading^31,46,48,49^. While spreading necessitates malleability, mechanical resilience is facilitated through properties such as tensile strength. Various studies have linked keratins to both these functions in culture cells and mouse embryos^31,50^; however, it remains unclear exactly how keratins could mediate these contrasting roles in a systemic manner. Our findings suggest a mechanism by which to reconcile the functions of keratins in ensuring tissue integrity^33,48,51^ and promoting tissue spreading^31,52^. The ability of keratins to resist mechanical stress increases with the progressive transformation of keratin organisation from an immature disconnected form to a dense, interconnected cellular network which finally organises into a supracellular tissue-scale network through desmosomal junctions between cells^22^. To ensure that this increasing resistance against deformation of the EVL does not stall EVL spreading and epiboly movements, keratins also facilitate tension-dependent actin network accumulation within the YSL, thereby adapting mechanical pulling force production within the YSL to EVL tissue viscosity resisting its deformation.

Keratins have previously been shown to interact with the actomyosin cytoskeleton^46,53^, although little is yet known about the biochemical realisation and functional significance of such interaction in early development. Keratins and actin are thought to interact directly^40,46^ or indirectly via large proteins, such as plakin and plectin^54,55^, and disruption of the actin network can interfere with keratin cytoskeletal organisation and stability^56^. Conversely, loss of keratin can affect actin network organisation during wound healing^50^ and actin stress fibre formation and cell polarisation in response to local tugging forces on C-cadherins^57^, pointing to a potential role of keratins in mediating force-dependent actin reorganisation. Our finding that keratins are required for actin mechanosensation within the YSL are consistent with these previous observations, providing direct evidence for keratins being involved in force-dependent actin network formation and contraction in multiple tissues, and mechanosensitive network maturation and relocation. How the keratin network functions in this process is not yet entirely clear, but is conceivable that it stabilises the actin network by providing a rigid substrate to which the actin network can couple.

Notably, keratin intermediate filaments are thought to be absent in insects^58^, suggesting that tissue morphogenesis and spreading in these animals can occur in the absence of keratin function. In embryogenesis of the insect *Tribolium castaneum*, for instance, the extraembryonic serosa, a simple squamous epithelial cell layer, undergoes massive spreading during epiboly^59^. Similar to EVL epiboly movements in zebrafish, serosa spreading is mediated by forces pulling on its leading edge^59^. Interestingly, this pulling leads to a regionalization of the serosa tissue into a solid-like dorsal portion with little cell rearrangements and a fluid-like ventral portion consisting of cells undergoing intercalations^59^. In contrast, no such clear regionalization can be observed in the zebrafish EVL with very little cell intercalations occurring throughout the tissue except some cells at the EVL margin withdrawing from the leading edge at very late stages of EVL epiboly. This different response of the EVL and serosa tissues to pulling forces might be due to the presence and absence of keratin expression within the respective tissues, pointing to the intriguing possibility that the function of keratins for homogeneous tissue spreading has become dispensable in insects. How this function of keratins in epithelial tissues has been adapted to specific organismal settings, and why keratins became expendable in many insect species remains to be explored.

While keratins belong to the most abundant and diverse cytoskeletal components in epithelial cells, remarkably little is yet known about the mechanisms by which they function in epithelial tissues. Our findings identify a critical role of keratins in promoting tissue viscosity and contractility in response to tissue tension. This ensures robust tissue spreading by balancing tissue integrity and expansion.

## Materials and methods

### Experimental Model and Subject Details

Zebrafish (*Danio rerio*) were maintained in the aquatics facility at ISTA, and embryos were collected, raised at 28-31°C and staged as previously described [Westerfield 2007] [Kimmel et. al. 1995]. The following wild-type (WT) and transgenic lines were used in this study: TL and AB wild-type and Tg*(krt18:Krt18GFP)*, Tg*(actb2:Utrophin-mcherry)*, Tg*(actb2:Lifeact- GFP),* and Tg*(acbt2:Utrophin-mcherry, krt18:Krt18GFP)* transgenic strains. All experiments were carried out according to the local regulations (Breeding 2023-0.288.351), and all procedures were approved by the Ethics Committee of IST Austria regulating animal care and usage.

### qPCR

Total RNA was extracted from 20 embryos at 1k (3.3 hpf), sphere (4 hpf), shield (6 hpf) and bud stage (9.5 hpf) using 750 µl Trizol as previously described. DNA-free™ DNA Removal Kit (Thermo Fisher Scientific) was used to clear the Genomic DNA following the manufacturer’s protocol. 3µg of RNA for each sample was taken as starting material to produce cDNA. NoRT control samples were produced with the Maxima H Minus First Strand cDNA Synthesis Kit following the manufacturer’s protocol. Linear amplification for the primers was tested using a series of dilutions for the cDNA to generate a standard curve. A 1:10 cDNA dilution was then chosen for the qPCR amplifications. For normalisation, the elongation factor 1 α (EF1α), as a housekeeping gene, was amplified (Miesfeld et al., 2015). The following primers for *keratin(krt)4, 5*, *8* and *18 were used:*

*krt4 forward: GCAGTCTATGAGGCTGAACTCC*
*krt4 reverse: CTCAGCCTTTGTTGAGCGGA*
*forwardkrt5 forward: ACT TCC TTC AAA ACC TTC AC*
*krt5 reverse: CCA GAT CCT GCT CCA AAA C*
*krt8 forward: CCA CCT ACA GCA AGA AAA CC*
*krt8 reverse: AGAGATGAAGCCACTACCAC*
*krt18 forward: GTAACATCCAGCATCAGACG*
*krt18 reverse: CACAACCTTTCCATCCACC*

The qPCR runs were performed on the Bio-Rad C1000 Thermal Cycler in triplicates using the Luna qPCR master mix.

### CRISPR/Cas9 mutant generation

To generate F0 crispants/mutants, Alt-R Cirspr-Cas9 kit was adapted to use a triple guide- mediate knockout of both keratin4 and keratin8 genes as described. For each guide RNA, 1μL crRNA 200μM; 1μL tracrRNA 200 μM were annealed in 1.5 μL Duplex buffer (IDT) by heating to 95°C for 5 mins and subsequent cooling on ice. 3 gRNAs were designed against 3 distinct exonst o target the whole locus of keratin4 and keratin8, each. RNPs were generated by annealing the gRNAs with Cas9 protein at 37°C for 15 mins. For keratin4/keratin8 mutants, 3 cas9 RNPs at 28.5 fmol (1000 pg) total gRNA together with 28.5 fmol (4700 pg) Cas9 protein (1 Cas9 to 1 gRNA) (IDT) against each gene were injected into one-cell stage embryos. The phenotype and survival rate were scored after each injection to test the efficacy of the knockouts after each injection.

### Cloning of expression constructs

Total RNA was extracted from 20 WT embryos at 4 and 8 hpf after dechorionation using 750 µl Trizol (Invitrogen). The cDNA library was generated with the Superscript III reverse transcription kit following the manufacturer’s instructions. The coding region of zebrafish *krt 18* was isolated using the following primers:

forward: 5’- GGGGACAAGTTTGTACAAAAAAGCAGGCTTAATGAGTCTGAGAACAAGCTACAGCG-3’ reverse: 5’- GGGGACCACTTTGTACAAAGAAAGCTGGGTTTTAAAGTTTCCTCTCCTTGGTTTCT GTGC -3’.

The dominant negative (DN) version of *krt18* was generated by mutating the Arginine to Cysteine at position 93 using the following primers:

forward: 5’-CATGCAGAACTTGAACGACTGTCTGGCCTCCTATCTGGAG -3’reverse: 5’-CTCCAGATAGGAGGCCAGACAGTCGTTCAAGTTCTGCATG -3’

The template DNA was then digested using dpn1 and the cDNA fragments were cloned into a pDEST plasmid using Gateway cloning. After transformation in NEB 5-alpha Ecoli strain in a pCS2 plasmid, the clones with the correct sequences were selected using sequencing.

For *krt 8* and *4*, the genomic fragments were isolated from cDNA libraries obtained from RNA of 8 hpf embryos as described above. The following primers were used to isolate specific DNA fragments: *krt8* forward: 5’- GCATGGACGAGCTGTACAAGAAGACAGAAAACACACAAGGCAGGATGAGTACG

-3’ *krt8* reverse: 5’- GCTGGTTTTCTTACTATACGTACTCATCCTGCCTTGTGTGTTTTCTGTCTTCTTG -3’ *krt4* forward: 5’- GGCATGGACGAGCTGTACAAGCTCAAAGACACGGGGATCATGTCGACGCGCTCTATCTCT -3’ *krt4* reverse: 5’- GTAATACGACTCACTATAGTTCTAGAGGCTTAATAGCGTTTACTGCTGACGGTGG -3’

The PCR products were then integrated to generate entry vectors via recombining with pDONR(P1-P2) (Lawson#208) and the entry clone was further recombined with pCS-N-term- mEmerald (Lawson #223) or pCS-N-term-mCherry (Lawson #362) destination vector (*krt4- mcherry, krt8-mcherrry, krt4-mEmerald, krt8-mEmerald*) or p3E mNeonGreen, pCS2-Dest (Lawson #444) for C-terminal tagging (*DNkrt18*).

### mRNA and *morpholino* injections

Constructs for obtaining *krt4*, *8*, and DN*krt*18 mRNA were generated as described above (cloning of expression constructs). mRNA was transcribed using the SP6 mMessage mMachine Kit (Ambion). For injections, glass capillaries (30-0020, Harvard Apparatus) were pulled using a needle puller (P-97, Sutter Instruments) and mounted on a microinjection system (PV820, World Precision Instruments). Injections at the one-cell stage were performed as previously described^60^. YSL injections were performed at high-stage (3.3 hpf) by injecting directly into the newly formed YSL through the yolk. 25pg of *krt*4*-mcherry* and *krt8-mcherry* mRNA were injected in one-cell stage embryos. Injecting higher mRNA amounts of single keratin isoforms (>100pg) led to aberrant keratin network formation. For lowering actomyosin contraction within the YSL, 100pg *CAMypt* mRNA together with 0.2% phenol red was injected into the YSL at high-stage (3.3 hpf). As controls, embryos were injected with 0.2% phenol red alone into the YSL at high-stage (3.3 hpf)^10,11^. For increasing actomyosin contraction within the YSL, 0.5-1pg *CARhoA* together with 2pg *H2A*-mCherry mRNA were injected into marginal blastomeres at the 128-cell stage as previously described^11^. Keratin intensity and density analyses were restricted to the regions within the YSL where mCherry labelled nuclei were clearly detectable.

For *morpholino* (MO) injections, *krt4, 8, and 18* MOs were designed as previously described^37^:

*krt4* MO: AGACCTGGTTGACATGATGCCTGTG

*krt8* MO: GGTTTTCTTGCTGTAGGTGGACATC

*krt18* MO: TGTAGCTTCTTCTCAGACTCATGGT

1ng of each MO together with 0.2% phenol red was injected either at the one-cell stage (for uniform knock-down) or at the high-stage (3.3. hpf) (for YSL-specific knock-down). As control, 2ng of a human *beta globin* MO (5′ - ‘CCTCTTACCTCAGTTACAATTTATA’ - 3′, Gene Tools) was injected.

### Embryo mounting and imaging

For inverted embryo imaging, embryos were dechorionated and mounted in 0.3%–0.5% low melting point (LMP) agarose in E3 (Invitrogen) on glass bottom dishes (MatTek) and then imaged on a Nikon CSU W1 with a CFI Plan Apo VC 60x WI/ NA 1.2 / WD 0.28-0.31 mm / Water, CFI Plan Apo λ 40x air / NA 0.95/ 0.17-0.25 mm or Leica Stellaris 8 (pipette aspiration experiments). For upright imaging, moulds were made using 3% agarose with wells in which the embryos were mounted in a lateral position and covered with 0.5%-0.6% low-melting point agarose. Samples were then imaged on a Leica SP5 or Leica SP8 microscope with a HC FUOTAR L 25x / 0.95 W (# 15506374), WD=2.5 mm, wide angle (41°) objective. Fixed samples were mounted in 0.5%–1% LMP agarose and put into prepared agarose moulds (2%) for upright imaging. Live embryos were imaged at 28.5°C ± 1°C.

### Analysis of EVL spreading

To determine EVL progression throughout development, the margin of the EVL was tracked along the orthogonal section of laterally imaged embryos, allowing to accurately track EVL movement over the yolk surface. The interval time was set to be 15mins for the aqusitions. In case of shorter acquisition time intervals, the EVL progression was interpolated to gain tracking intervals of 15 min. The speed measured by this method was identical with the speed measured by PIV flows described below (analysis of actomyosin flows).

### UV cell ablations

Nikon CSU W1-01 SoRa+NIR spinning disk microscope with a home-built UV laser ablation system was used for ablating cells and monitoring the wound healing response. Ablations were performed by a single UV-laser ablation line over at least 4-5 cells with the intensity of the UV laser set at a level causing spontaneous cell delaminations. Z-stacks of 5-10 μm were acquired at 30 sec intervals. EVL cells were segmented with Cellpose^61,62^ as described below (segmentation and tracking of EVL movements) and the Feret angle (angle of the longest axis) was measured using Fiji. With the wound set as the centre, the polar coordinates of each cell were calculated and the angle with respect to the polar angle was calculated by taking the absolute difference of the polar angle and the Feret angle. 3 radial bins of 600 pixels were taken to estimate the average alignment measured. For a perfectly rosetted aligned structure the alignment measured would be 0. All angles were transformed to be between 90° and -90°

### Pipette aspirations

Pipette aspirations were performed to measure the viscosity of the EVL as previously described. In short, pipette aspirations were carried out on embryos mounted in 3% methylcellulose in E3 on an inverted Leica SP5 or a Leica Stellaris 5 confocal microscope equipped with the micropipette aspiration system. For creep recovery experiments to measure viscosity, fire-polished and with heat-inactivated FBS passivated micropipettes (Biomedical Instruments) with an inner diameter of 60 μm, 30° bent positioned by micromanipulators (TransferMan Nk2, Eppendorf), and with a blunt end were placed on the EVL around 3-4 cells away from the EVL margin. A negative pressure was applied with an increment of 10 Pa using a Microfluidic Flow Control System Pump (Fluiwell, Fluigent) and Dikeria micromanipulation software^63^. Images were acquired every sec. A negative pressure of 200 mbar was applied until the aspirated tissue flowed in the pipette with constant velocity, followed by pressure release. The length of the aspirated tissue was measured using a custom-made Fiji macro and plotted over time to calculate the aspiration and retraction speed. The viscosity was calculated as previously described^64,65^. Pipette aspirations were also used to study the mechanosensitive response of the keratin cytoskeleton within the EVL and the actin cytoskeleton within the YSL. For the EVL aspirations, the EVL of Tg*(acbt2:Utrophin-mcherry,krt18:Krt18GFP)* or Tg*(kert18:Krt18GFP)* embryos was aspirated under constant pressure, and the mean intensity of keratin at the midplane of the pipette and a region of 30 µm around the aspiration site were recorded. For the YSL aspirations, a z-stack of 50 µm was taken and the actin intensity in Tg*(acbt2:Utrophin-mcherry,krt18:Krt18GFP)* embryos was measured on the YSL surface in intervals of 30 sec.

### Analysis of YSL actomyosin flows

To measure actin flow velocities within the YSL, Tg*(actb2:Utrophin-mcherry)* or Tg*(actb2: Lifeact-GFP)* embryos were mounted on a Nikon CSU W1 spinning disc microscope in 0.5% LMP agarose and imaged every 30 sec. A 500 × 500-pixel region of interest (ROI) within the YSL was selected with maximum intensity projections centred along the AV axis close to the margin of the EVL. The respective ROI was then used for PIV analysis using PIVlab^66^ and post-processed by a custom-made Python script. Flow velocities represented in the kymographs were averaged over three consecutive time points. Alignment of these flows was analysed by measuring the angular mean of the flow vectors in the ROI as described above.

### Segmentation and tracking of EVL cells

To visualize EVL cells, Tg*(actb2: Utrophin-mcherry) or* Tg*(acbt2:Utrophin-mcherry, krt18:Krt18GFP)*transgenic embryos were used to mark apical cell junctions. Maximum intensity projections of Z-stacks covering approximately 150 µm depth (for movies acquired with a SP8 confocal microscope) or 30 µm depth (for movies acquired on a Nikon spinning disc microscope) were generated for subsequent analysis. Segmentation of the EVL cells was performed using a "human in the loop" pipeline using the Cellpose segmentation software. Initial segmentations were generated using models trained on manually annotated datasets. These models were then employed to make further segmentations. Post-processing involved manually correcting any segmentation errors and using these corrections to further refine the segmentation models. Heightmaps were generated with the Local Z-projector plug-in in Fiji in order to correct for the curvature of the embryo. Deproj functions were used to integrate the segmentations and the heightmaps to measure EVL cell area and alignment. Data were processed and plotted using custom python scripts.

### Keratin network segmentation

For measuring the network of the keratin cytoskeleton in EVL cells, Tg*(krt18:Krt18GFP)* embryos were imaged at 63x magnification as described above (embryo mounting and imaging). As most of the keratin network within EVL cells formed a single sheet at the apical surface, the Z-projections of the keratin network over the entire EVL cell were used. First, a Gaussian filter and denoising were applied to obtain the filamentous skeleton. Next, the images were thresholded using an adaptive median threshold and the area fraction covered by the network as an estimate of network density was determined using Fiji. For junctional keratin accumulation, the cell margins were outlined using EVL segmentations as described above (segmentation and tracking of EVL cells) with a thickness of about three pixels to obtain a skeleton junction image. The skeleton junction image was then used to segment the junctional pool of keratin in each cell and the intensity of the junctional pool was measured in the segmented image. The total intensity of keratin per cell was measured using the whole apical segmented area inside the junctional segmentations.

### Fluorescence in situ hybridization

WT or Tg*(krt18:Krt18GFP)* embryos at 9hpf were fixed in 2% paraformaldehyde at RT for an hour and stepwise fixation with and subsequent storage in methanol at -20°C. After overnight fixation in methanol the embryos were stepwise rehydrated to PBST (phosphate buffered saline, 0.1% Tween-20 pH 7.3) followed by incubation in hybridization buffer (10% Dextran sulfate, Formamide 10%, tRNA 1mg/ml, SSC 2x, BSA 0.02%, Vanadyl-ribonucleoside complex (NEB S1402S) 2mM) at 30°C. Stellaris RNA FISH probes for krt8 mRNA were designed from zebrafish mRNA sequences using LGC Biosearch Technologies’ Stellaris® RNA FISH Probe Designer and 30 probes per probeset were ordered. The embryos were incubated in this probset overnight at 30°C. The hybridized embryos were washed with Wash buffer 3 times before mounting in gold mounting media before imaging at 63x as described above.

### EVL primary cell culture

At 3.5 hpf or at 6 hpf, Tg*(acbt2:Utrophin-mcherry,krt18:Krt18GFP)* or Tg*(krt18:Krt18GFP)* embryos were transferred to pre-warmed (28.5–31°C) 0.9x DMEM/F12 medium supplemented with GlutaMAX and Penicillin-Streptavidin as described (ref). The blastoderm caps were dissected from the yolk cells at sphere stage with forceps and transferred to 1.5 ml Eppendorf tubes using glass pipettes. These explants were dissociated by gentle tapping and seeded on bilayers at 29°C and imaged for at least 4 hours after seeding. EVL cells were identified by their shape and the presence of keratin fluorescence.

---

**Figure supplement 1:**
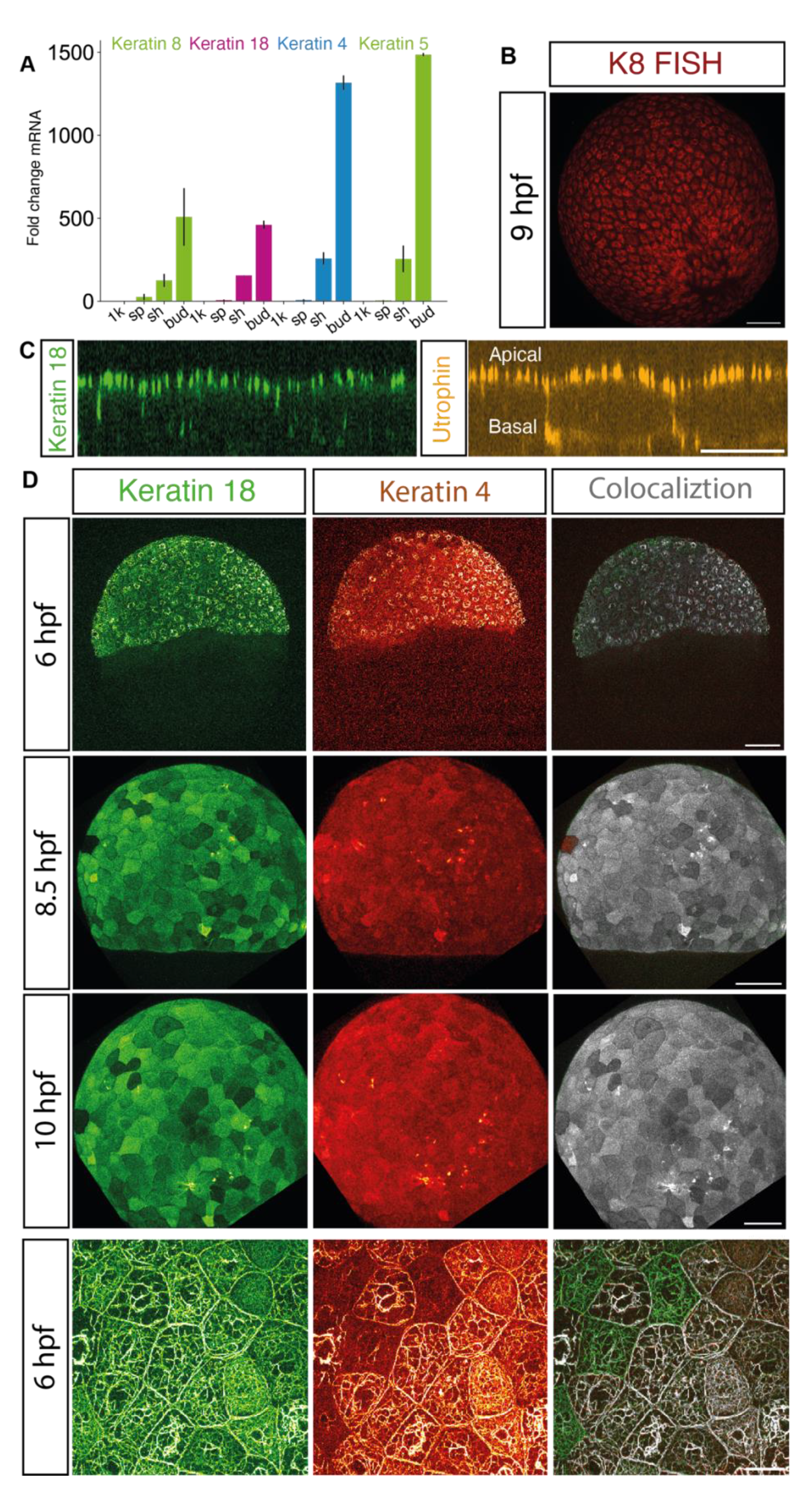
keratin expression and localization during epiboly. (A) Bar plot of fold change of *keratin 18*, *keratin 8*, *keratin 4*, and *keratin 5* expression in embryos at 1K (3.3hpf), 30% epiboly (4.5 hpf), shield (6hpf) and bud (9hpf) stages measured by qPCR. (N=3, n=46 embryos; p-values k8: 0.4833, k18: 0.043 ANNOVA). (B) Maximum Intensity projections of krt8 mRNA fluorescence via in situ hybridization in WT embryos at 9 hpf. Scale bar: 25 µm (C) Maximum Intensity projections of keratin network in Tg(*krt18: Krt18GFP)* embryos at shield (6hpf), 75% epiboly (8.5 hpf) and bud (10 hpf) stages injected of 50pg Krt4-mcherry RNA at the one-cell stage. Left column, keratin 18 (green); middle column, keratin 4 (red); right column, keratin 4 and 18 colocalization (white; co-localization index R below threshold 0.005). Scale bar: 100 µm (D) Maximum Intensity projections of keratin network in Tg*(krt18: Krt18GFP)* embryos at shield (6hpf), 75% epiboly (8.5 hpf) and bud (10 hpf) stages injected of 50pg Krt4-mcherry RNA at the one-cell stage. Scale bar: 25 µm

**Figure supplement 2:**
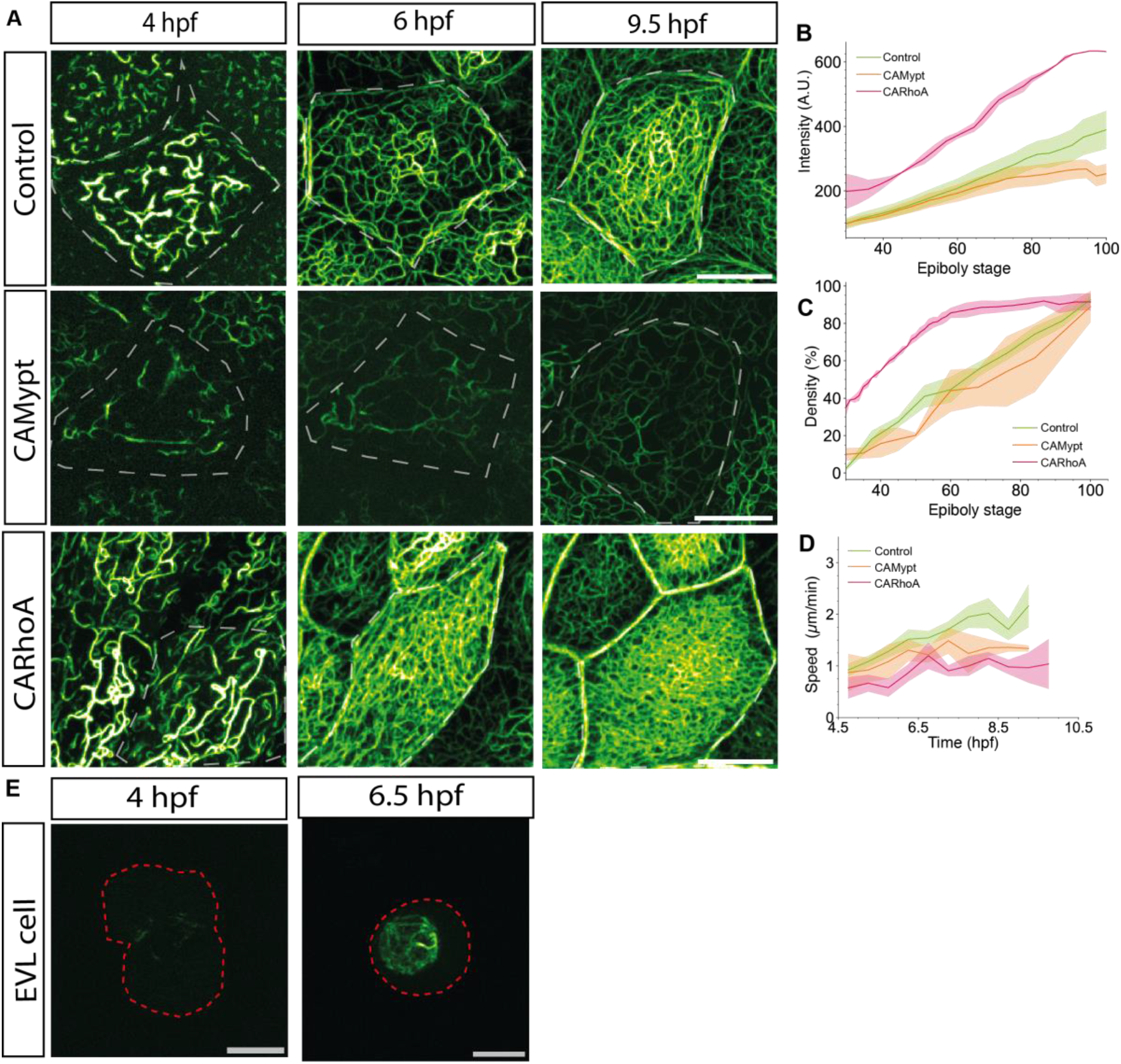
Tension-dependent regulation of keratin expression within the EVL. (A) Maximum intensity projection images of keratin expression in Tg(*actb2: Utrophin-mcherry, krt18:Krt18GFP)* embryos at sphere (4 hpf, left column), shield (6 hpf, right column), and 90% epiboly (9.5hpf, right column) stages injected with (0.2% phenol red, control, top row), 100pg *caMypt* (middle row), or 50 pg *caRhoA* (bottom row) directly into the YSL at 3.3 hpf (control, *caMypt*) or into marginal cells at 3 hpf (*caRhoA*) Scale bar: 25 µm (B) Plot of average keratin intensity as a function of epiboly stages in Tg(*krt18:KrtGFP)* control (green, N=3, n= 5 embryos), *caMypt* (orange, n=3, n=5 embryos) and *caRhoA* mRNA injected embryos(pink, N=3, n=4 embryos) at 3.3 hpf into the YSL as described in Figure 2(A- C). Error bars as ribbons SD of mean. (C) Plot of average density of keratin network as a function of epiboly stages in Tg(*actb2:Utrophin-mcherry, krt18:Krt18-GFP)* control (green, N=3, n= 6 embryos), *caMypt* (orange, n=3, n=4 embryos) and *caRhoA* mRNA injected embryos (pink, N=3, n=3 embryos) at 3.3 hpf into the YSL as described in (B). Error bars as ribbon SD of mean of individual cells per replicate. (D) Plot of EVL epiboly movement speed as a function of time (hpf) during epiboly starting at sphere stage (4hpf) until late epiboly stages (9 hpf) in Tg(*actb2: Utrophin-mcherry, krt18:Krt18GFP)* embryos injected at 3.3hpf with control (green, N=3, n= 6 embryos), *caMypt* (orange, n=3, n=4 embryos) and *caRhoA* mRNA (pink, N=3, n=3 embryos). Error bars as ribbon SD of mean of individual cells per replicate. (E) Maximum intensity projection images of keratin expression in individual primary cells from Tg(*krt18:Krt18GFP)* embryos dissociated at 4 hpf (left) and 6.5 hpf (right). The outline of the cell is shown as the red dotted line. Scale bar: 15 µm

**Figures supplement 3:**
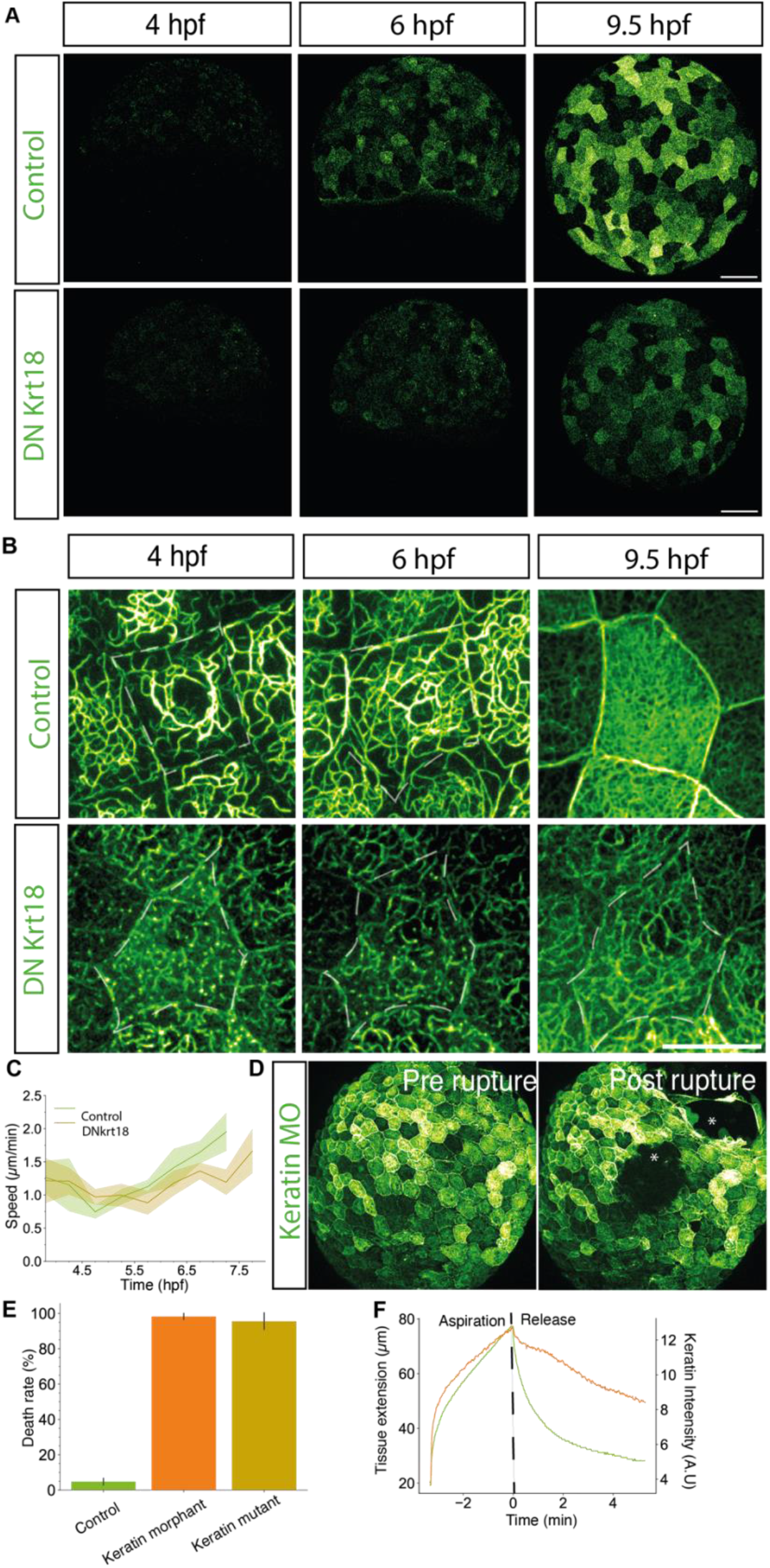
Effect of dominant negative keratin 18 expression on keratin network formation and EVL epiboly. (A) Maximum intensity projection images of keratin expression in Tg(*actb2: Utrophin-mcherry, krt18:Krt18GFP)* at sphere (4hpf, left column), shield (6phf, middle column) and bud (10 hpf, right column) stages injected with 0.2% phenol red (control, top row) or 150 pg dnKrt18 RNA (bottom row) at one-cell stage. Scale bar: 100 µm (B) Maximum intensity projection images of keratin expression in Tg(*actb2: Utrophin-mcherry, krt18:Krt18GFP)* at sphere (4hpf, left column), shield (6phf, middle column) and bud (10 hpf, right column) stages injected with 0.2% phenol red (control, top row) or 150 pg dnKrt18 RNA (bottom row) at one-cell stage. Scale bar: 25 µm (C) Plot of EVL epiboly movement speed as a function of time (hpf) during epiboly starting at sphere (4 hpf) until late epiboly (9 hpf) stages in Tg(*actb2: Utrophin-mcherry, krt18:Krt18GFP)* embryos injected with 0.2% phenol red (control, green) or 150 pg dnKrt18 RNA (orange) at one-cell stage. (N=2, n=6 embryos). (D) Maximum intensity projection images of EVL during rupture at 13.5 hpf in Tg(*actb2: Utrophin-mcherry, krt18:Krt18GFP) embryos injected with* 1ng *krt4* plus 1ng *krt8* MO at one cell stage showing the EVL before (top) and after the EVL ruptures (bottom). (E) Bar plot of quantification of death rate in Tg(*actb2:Utrophinmcherry, krt18:Krt18GFP)* embryos injected at the one-cell stage with 2ng control MO (top row; green, N=4, n=4 embryos), 1ng *krt4* plus 1ng *krt8* MO (middle row; orange, N=4, n=4 embryos), or with TraCr *krt4* and *krt8* gRNA (*krt4/8* crispant F0; bottom row; yellow, N=2, n=6 embryos). (F) Exemplary plot of tissue extension as a function of EVL tissue aspiration time, displaying a linear response upon aspiration and retraction(green). Plot of fluorescence intensity of keratin as a function of aspiration in the same embryo measured in the medial plane of the pipette in the same embryo (orange)

**Figures supplement 4:**
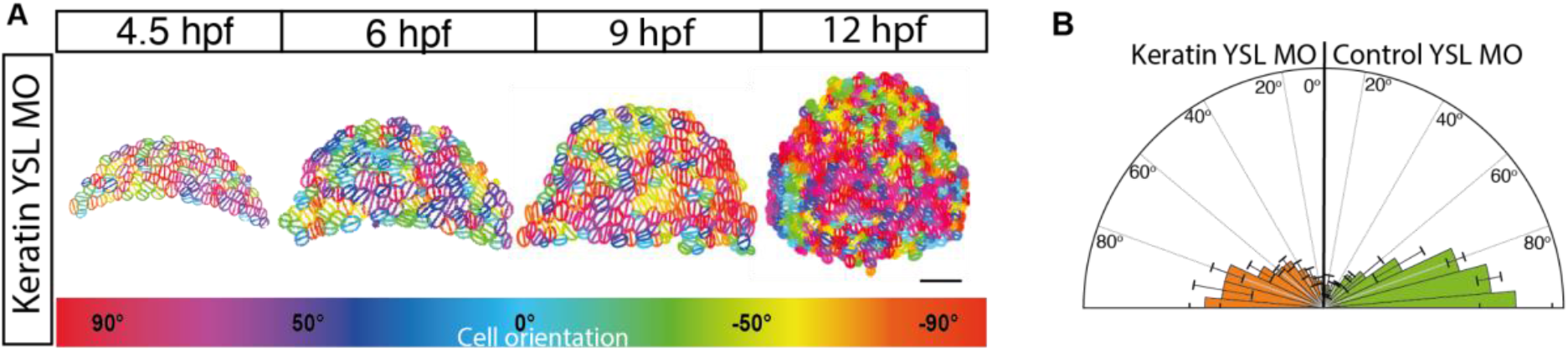
EVL cell alignment along the axis of tissue spreading is independent of keratin expression within the YSL. (A) Exemplary plots of EVL cell orientations with ellipses representing shape descriptors (long and short axis) of individual EVL cells with the line in the middle marking the orientation of the long axis at consecutive stages during epiboly (4.5, 6, 9, 12 hpf) in curvature corrected Tg(*actb2:Utrophin-mcherry, krt18:Krt18GFP)* embryos injected at high stage (3.3 hpf) directly into the YSL to interfere with keratin network formation within the YSL specifically. Each cell is colour-coded according to the orientation of the axis (hsv) as shown in the colour bar at the bottom (Red: AV axis orientation, blue: dorsoventral/DV orientation). (B) Rose plot of EVL cell orientations in curvature corrected *Tg(actb2:Utrophinmcherry,krt18:Krt18GFP)* embryos at 6 hpf injected at high stage (3.3 hpf) into the YSL with 2 ng control MO (B; N=4, n=93 cells) or 1ng *krt4* plus 1ng *krt8* MO (C; N=4 n= 84 cells).

**Figures supplement 5:**
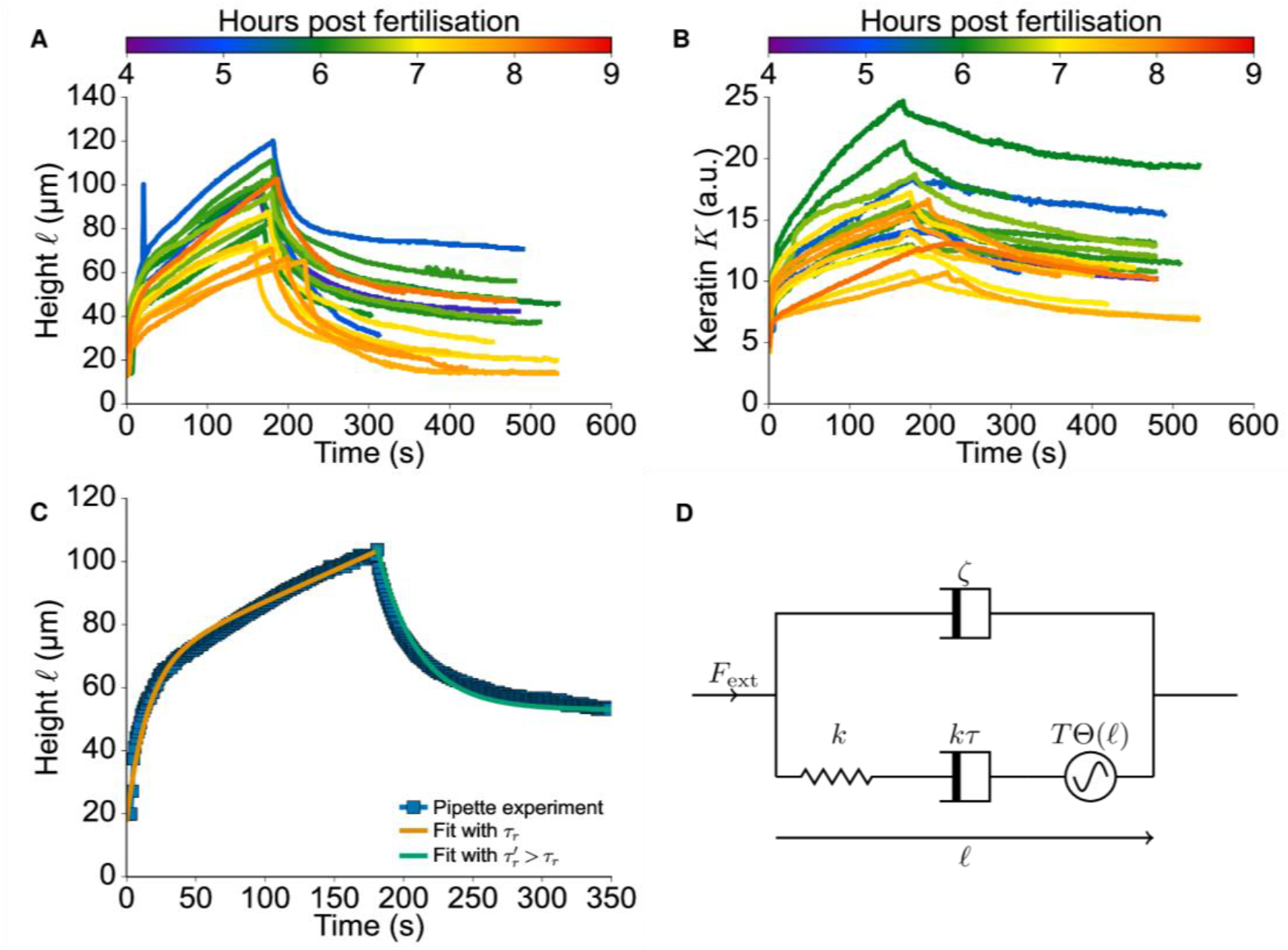
Micropipette aspiration experimental data and Maxwell model. (A) Plot of the height of the aspirated EVL tissue as a function of time upon micropipette aspiration. Colormap represents developmental time starting at the beginning of epiboly (4hpf, blue) to later stages (8 hpf, orange). (B) Plot of keratin expression as a function of time upon EVL micropipette aspiration at the center of the pipette in a single Z-slice. Colormap represents developmental time starting at the beginning of epiboly (4hpf, blue) to late stages (8 hpf, orange). (C) Representative plot of the height of the aspirated EVL tissues as a function of time (blue, square, spline) upon micropipette aspiration fitted to the functional form equation (S4) to separately obtain the relaxation times during aspiration (𝜏_𝑟_, yellow) and release ( 𝜏_𝑟_*′*, green). (D) Diagram of the Maxwell model coupled to substrate friction (equation (S1)), constructed to infer parameters from the aspiration experiments. See SI section 1 for details.

**Figures supplement 6:**
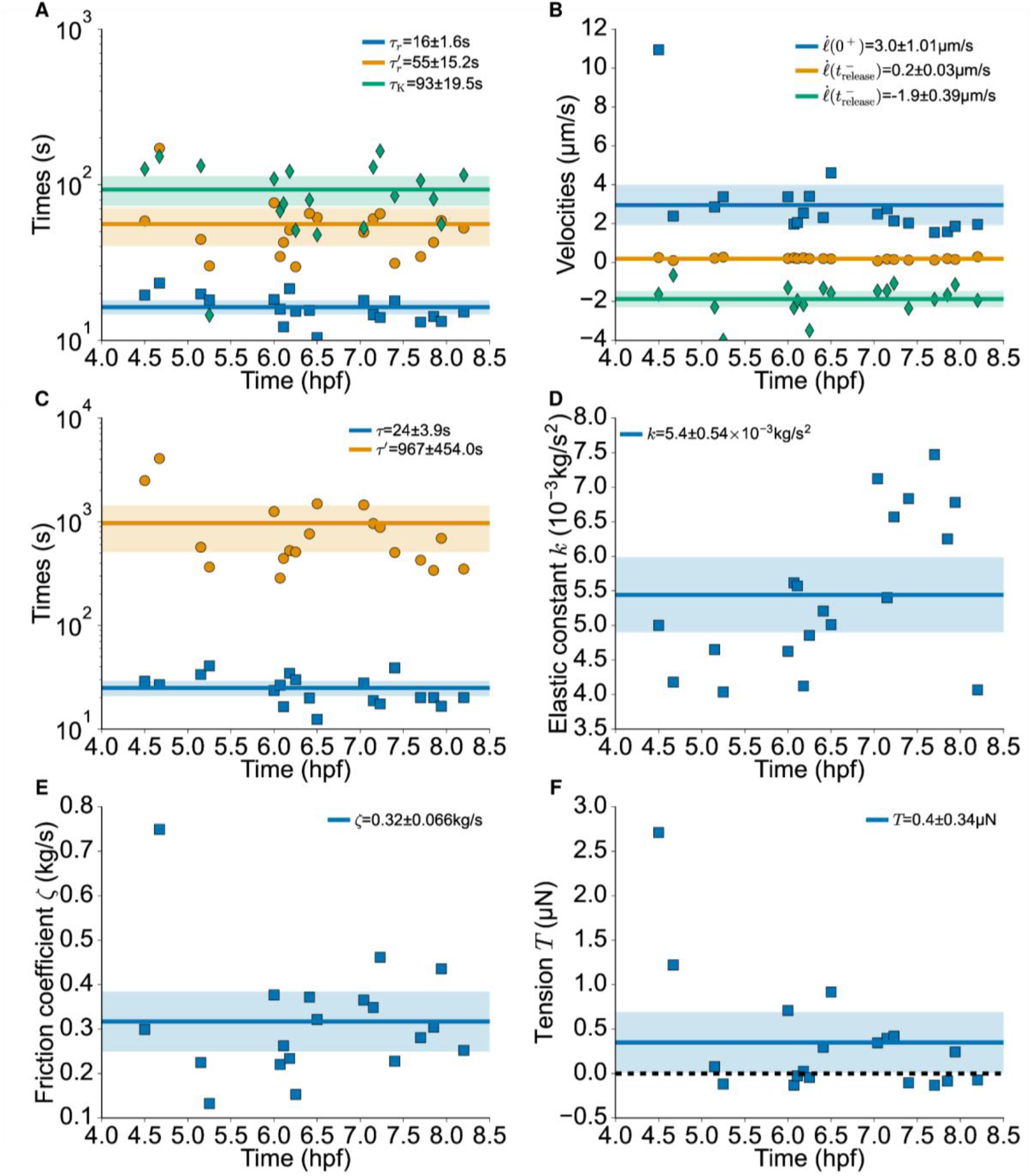
Time and velocity scales for the EVL micropipette aspiration experiment and parameters of the Maxwell model. (A) Scatter plot of measured time scales 𝜏_𝑟_ (blue, square) and 𝜏_𝑟_*′* (green, rhombus) in the EVL aspiration experiments from fits of the height relaxation to equation (S4) during the aspiration and release phases, respectively, using the method described in figure supplement 5C. Measured time scale 𝜏_𝐾_ for the relaxation of the keratin K(T) as depicted in (figure supplement 5B) after pipette release (yellow, circle), from a fit to exponential form 𝑒^−𝑇/𝜏𝐾^. Line plots depict the average with confidence intervals for 𝜏_𝑟_ (blue), 𝜏_𝑟_ *′* (green) and 𝜏_𝐾_ (yellow). (B) Scatter plot of measured velocities in the EVL aspiration experiments as extracted by linear fits to the height curves ℓ(t) (figure supplement 5A) where 𝑡_𝑟𝑒𝑙𝑒𝑎𝑠𝑒_ is the time where the pressure is removed. (C-F) Fits to equation (S4) of the EVL aspiration experiments during aspiration and release inverted to extract Maxwell model parameters (equations (S5)), i.e. tissue relaxation time scales 𝜏 during aspiration and 𝜏*′* during release (C), tissue elastic constant k (D), substrate friction coefficient ζ (E), and internal tissue tension T (F). Line plots over the scatter are the average with confidence intervals shaded. See SI section 1 for model details.

**Figures supplement 7:**
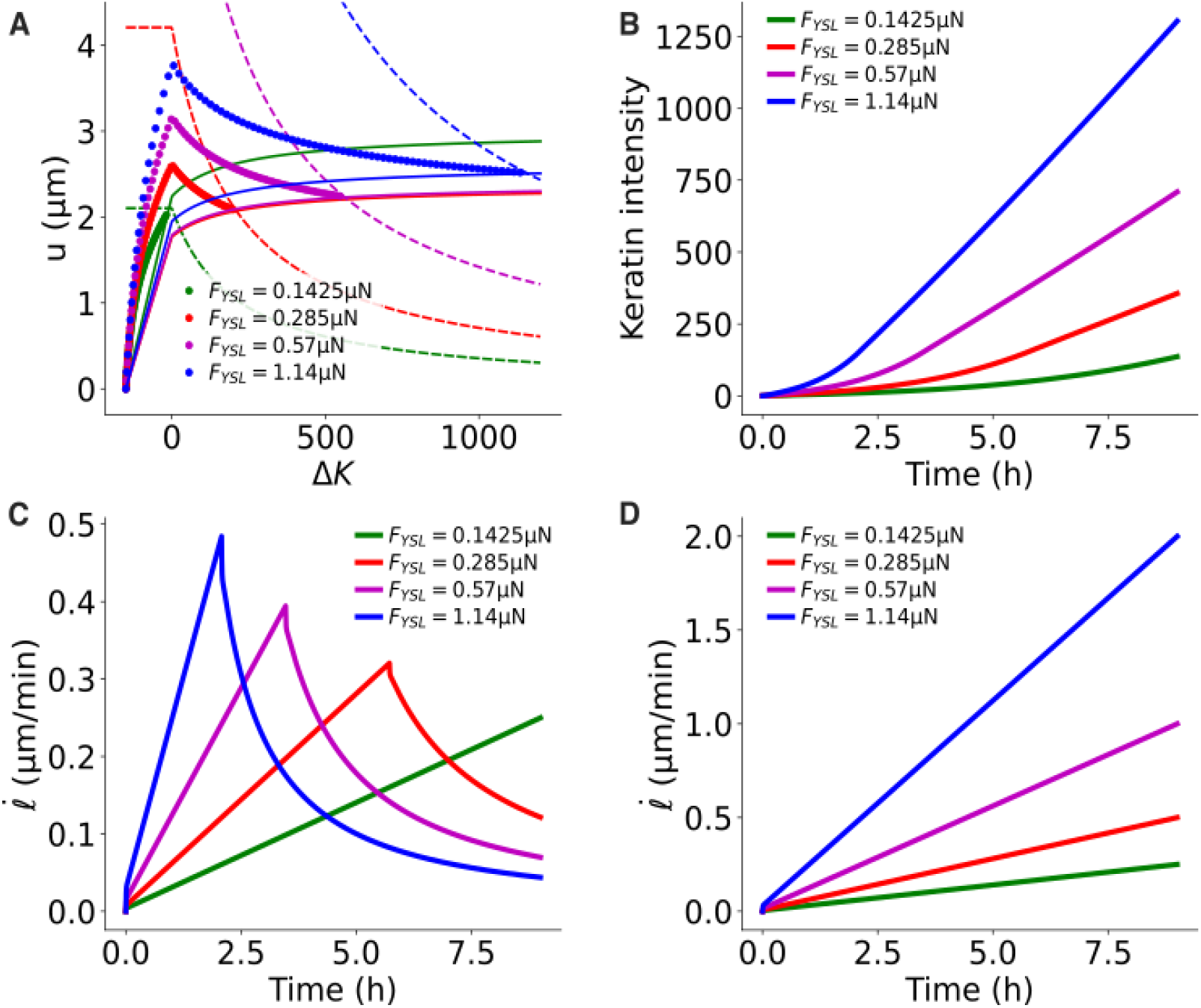
Mean-field dynamics of simulated EVL model tissue. (A) Reduced variable phase space (𝐾 − 𝐾_𝑡_*_ℎ_*, 𝑙 − 𝑙*_0_*) in EVL tissue models for different values of 𝐹_𝑌𝑆𝐿_ (linear force ramp). Simulations with feedback (𝛽 = *0*.*005*) are shown as dots, together with the nullclines of 𝐾 − 𝐾_𝑡_*_ℎ_* (solid) and of 𝑙 − 𝑙*_0_* (dashed). (B,C) Keratin dynamics (B) and junction length expansion speed *i*(C) as a function of time during EVL tissue model expansion. (D) Junction length expansion speed *i* for the keratin deficient EVL tissue model with no feedback (𝛽 = *0*). See SI section 4 for model details.

## Video Legends

Video 1: Keratin expression within the EVL during epiboly

Time-lapse of keratin expression (right) and network organization (left) in Tg*(krt18:Krt18GFP)* embryos during epiboly (4 - 9.5 hpf). Lateral view of embryos imaged at 11.25 mins/frame. Scale bar, 100 (left) and 30 µm (right).

Video 2: Keratin expression within the EVL in embryos with reduced YSL pulling force

Time-lapse of keratin expression within the EVL upon mechanical pulling force reduction in a representative Tg*(krt18:Krt18-GFP)* control embryo (YSL injection of 0.2% phenol red, left) and an embryo injected with 100pg *CAMypt* into the YSL (right). Lateral view of the embryos imaged at 10 mins/frame from 4.5-11 hpf. Scale bar, 100 µm.

Video 3: Keratin network maturation in EVL cells of embryos with reduced YSL pulling force

Time-lapse of keratin network maturation in EVL cells upon mechanical pulling force reduction in a representative Tg*(krt18:Krt18-GFP)* control embryo (YSL injection of 0.2% phenol red, left) and an embryo injected with 100pg *CAMypt* into the YSL (right). Lateral view near the EVL/YSL margin imaged at 10 mins/frame from 4.5-7.1 hpf. Scale bar, 25 µm.

Video 4: Keratin expression within the EVL of embryos with enhanced YSL pulling force

Time-lapse of keratin expression upon mechanical pulling force increase in representative Tg*(krt18:Krt18-GFP)* control embryo (YSL injection of 0.2% phenol red, left) and 50 pg *CARhoA* (marginal cell injection at 3.3 hpf, right). Lateral view imaged at 10 mins/frame from 4.5-10.3 hpf. Scale bar, 100 µm.

Video 5: Keratin network maturation in EVL cells of embryos with enhanced YSL pulling force

Time-lapse of keratin network maturation upon mechanical pulling force increase in a representative Tg*(krt18:Krt18-GFP)* control embryo (YSL injection of 0.2% phenol red, left) and and embryo injected with 50 pg *CARhoA* in marginal blastomeres at 3.3 hpf (right). Lateral view near the EVL/YSL margin imaged at 10 mins/frame from 4.5-7.5 hpf. Scale bar, 25 µm.

Video 6: Keratin expression within the EVL of embryos with reduced keratin type II expression

Time-lapse of keratin expression in representative Tg*(krt18:Krt18-GFP)* embryos injected at the one-cell stage either with 2ng control MO (control, left) or 1ng *krt4* plus 1ng *krt8* MO (right). Lateral view imaged at 10.25 mins/frame from 4-15.75 hpf. Points of rupture in keratin morphant embryos are marked with asteriks. Scale bar, 100 µm.

Video 7: Keratin network maturation in EVL cells of embryos with reduced keratin type II expression

Time-lapse of keratin expression in representative Tg*(krt18:Krt18-GFP)* embryos injected at the one-cell stage either with 2ng control MO (control, left) or 1ng *krt4* plus 1ng *krt8* MO (right). Lateral view of the embryos imaged at 10 min/frame from 4.5-8.4 hpf. Scale bar, 25 µm.

Video 8: Changes in keratin expression upon EVL aspiration

Time lapse of EVL aspiration in a representative Tg*(krt18:Krt18-GFP)* embryo using a 60 µm pipette imaged by brightfield (left) and confocal (keratin, green, right) microscopy. Z-plane in the center of the pipette imaged at 1 sec/frame. Scale bar, 25 µm.

Video 9: Changes in keratin expression during EVL wound closure after cell ablation

Time lapse of EVL in a representative Tg*(actb2:Utrophin-mcherry, krt18:Krt18-GFP)* embryo imaged before (pre) and after (post) UV laser-mediated cell ablation showing keratin (right, green) and actin (left, orange). Imaged at 20 sec/frame. Scale bar, 25 µm.

Video 10: Wound closure in control and keratin deficient embryos

Time lapse of EVL response in representative Tg*(actb2:Utrophin-mcherry)* embryo imaged before (pre) and after (post) UV laser-mediated cell ablation injected at the one-cell stage either with 2ng control MO (control, left) or 1ng *krt4* plus 1ng *krt8* MO (right). Frame rate 20 sec/frame. Scale bar, 25 µm.

