## Supplementary material for "Keratins coordinate tissue spreading by balancing spreading forces with tissue material properties": SI

#### 1 Experimental measurements and visco-elastic Maxwell model

Pipette aspiration experiments are performed on the gastrulating tissue as follows: The tissue is initially at rest. At time $t = 0$ a constant pressure differential ($\Delta P_{\mathrm{ext}} = 2\,\mathrm{mbar}$) is applied within the pipette (diameter $d = 60\,\mu\mathrm{m}$), leading to a constant pulling force $F^0_{\mathrm{ext}} = (\pi/4)\,\Delta P_{\mathrm{ext}}\,d^2 \approx 0.57\,\mu\mathrm{N}$, causing the tissue to move upwards. Then at a later time, the pressure differential is removed, i.e. $\Delta P_{\mathrm{ext}} = 0$, causing the tissue to retract and move downwards in the second phase.

Figure S1A-B shows the height of the aspirated tissue and the corresponding keratin expression for tissues at different times post fertilisation. Tissues show viscous behaviour at long times (deformation at constant velocity).

During our pipette experiment approximately 5 cells are aspirated into the pipette, and move with a typical velocity in the 0.1-1 $\mu\mathrm{m\,s^{-1}}$ range. The standard model of inferring tissue viscoelasticity from pipette aspiration Ref. [64, 65] involves approximately 100 cells at a typical velocity of 0.01 $\mu\mathrm{m\,s^{-1}}$. We will therefore in contrast assume that our experiment probes the mechanical behaviour of *individual cells* rather than the whole tissue – it is consistent with junction cutting experiments which probe cell-level elasticity and report velocities in the 0.2-0.5 $\mu\mathrm{m\,s^{-1}}$ range [10]. Epiboly however involves $\gtrsim 500$ cells [13] with a typical velocity in the 0.01-0.03 $\mu\mathrm{m\,s^{-1}}$ range [10], and so we will use our results here to set the *individual cell* parameters for a vertex model of epiboly (see section 2).

##### Modified Maxwell model

To account for these observations, we introduce a simple model for a stressed viscoelastic tissue, namely a Maxwell model with additional internal tension, coupled with friction on a substrate (Figure S1D). Let $\ell$ be the length of the material, which we identify with the height of the aspirated tissue in the pipette experiment. We write the constitutive equations

$$\zeta\dot{\ell} = -k(\ell - \ell_0) - T\Theta(l) + F_{\mathrm{ext}}, \qquad \triangleright \text{ overdamped harmonic oscillator} \tag{S.1a}$$

$$\dot{\ell}_0 = \frac{1}{\tau}(\ell - \ell_0), \qquad \triangleright \text{ rest length relaxation} \tag{S.1b}$$

where $\zeta$ is the substrate friction coefficient, $k$ is the elastic constant of the tissue, $\tau$ its internal relaxation time of the tissue, $T$ is the internal tension in the tissue, $\Theta$ is the Heavisde function introduced to avoid unphysical $\ell < 0$, and $F_{\mathrm{ext}}$ the external applied force from the pipette. These lead to the following equation of motion for length $\ell$

$$\ddot{\ell} + \frac{1}{\tau_r}\dot{\ell} = \frac{1}{\zeta\tau}\left(F_{\mathrm{ext}} - T\Theta(l)\right) + \frac{1}{\zeta}\left(\dot{F}_{\mathrm{ext}} - T\frac{\mathrm{d}}{\mathrm{d}t}\Theta(l)\right), \tag{S.2a}$$

$$\tau_r = \frac{\zeta\tau}{\zeta + k\tau} < \tau. \tag{S.2b}$$

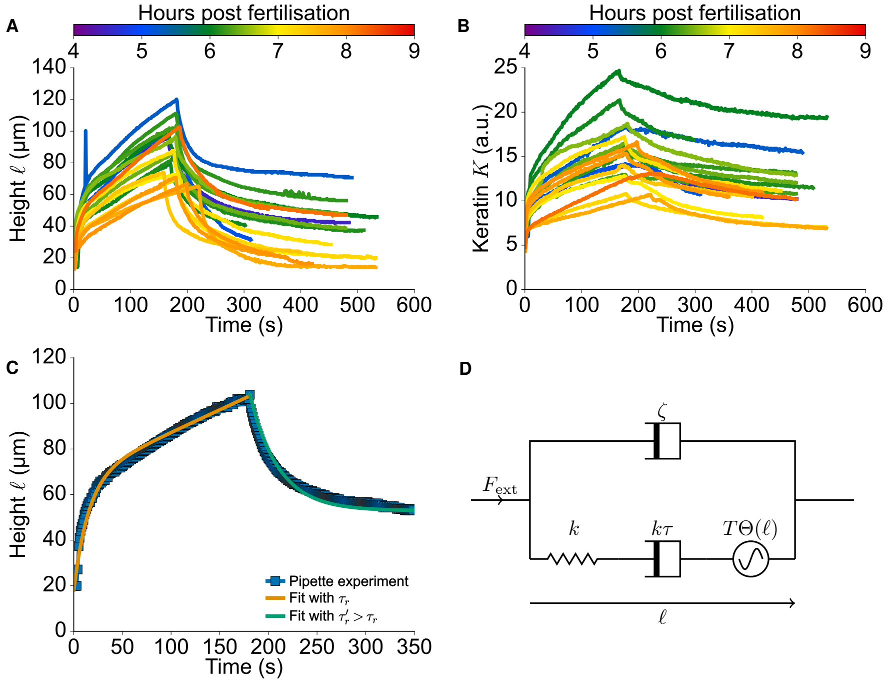


Figure S1: (A) Height of the aspirated tissue as function of time in a normal tissue with the corresponding (B) keratin expression level in arbitrary units. (C) Fit of a sample experimental curve to the functional form equation (S.4). (D) Diagram of the Maxwell model coupled to substrate friction.

Consistent with the pipette experiment, we consider $F_{\text{ext}}$ to be a piecewise constant function. Assuming that $F_{\text{ext}}$ and/or $\Theta(l)$ are discontinuous at time $t$, we compute the inital condition to solve (S.2a) at times $t' > t$ [67]

$$\ell(t^+) - \ell(t^-) = 0, \tag{S.3a}$$

$$\dot{\ell}(t^+) - \dot{\ell}(t^-) = \frac{1}{\zeta}\Big([F_{\text{ext}}(t^+) - F_{\text{ext}}(t^-)] - T[\Theta(\ell(t^+)) - \Theta(\ell(t^-))]\Big), \tag{S.3b}$$

and the form of the solution under these assumptions is

$$\ell(t) = (\tau_r(\dot{\ell}(0^+) - v_\infty) + \ell(0^+))\left[1 - \frac{\tau_r(\dot{\ell}(0^+) - v_\infty)}{\tau_r(\dot{\ell}(0^+) - v_\infty) + \ell(0^+)} e^{-t/\tau_r}\right] + v_\infty t, \tag{S.4a}$$

$$v_\infty = \frac{F_{\text{ext}} - T}{\zeta + k\tau}, \tag{S.4b}$$

provided that $\ell(t > 0) > 0$. Note that this form of the solution is identical to Ref. [64] but leads to different continuity relations, i.e. our model guarantees a continuous height function. Our model allows for a straightforward parametrisation of the internal elasticity and viscous relaxation through parameters $k$ and $\tau$, and the substrate dissipation through parameter $\zeta$, thus simplifying the link to the vertex model of section 2. We provide an mapping between the parameters of the two models in equations (S.6).

We now have four parameters ($\zeta$, $k$, $\tau$, and $T$) which could have time-dependent values because of the internal effect of increasing levels of keratin. We simplify the fitting problem by assuming that $\zeta$, $k$, and $T$ are constant on the timescale corresponding to the pipette experiment, leaving only $\tau$ variable. The effective relaxation time scale $\tau_r$ (S.2b) characterises how the velocity responds to the externally applied force. In the pipette experiment, we expect the effect of keratin to be stronger during the second phase (release) than during the first phase (aspiration). Therefore, we separately fit both phases to functions of the form (S.4) and extract the corresponding relaxation time scales $\tau_r$ and $\tau_r'$ in the first and second phases respectively (see Figure S1C). Figure S2A shows $\tau_r' > \tau_r$ by a factor $\approx 2$. Possible interpretations may be that the effect of keratin is to decrease the elastic constant $k$, which is counter-intuitive, or that it increases the relaxation time scale $\tau$ (*i.e.* the effective friction coefficient $k\tau$). We will consider the second case. Consequently we formulate these final hypotheses:

1. the relaxation time in the aspiration phase $\tau_r = \zeta\tau/(\zeta + k\tau)$ is related to a first internal relaxation time $\tau$,

2. the velocity before release $\dot{\ell}(t^-_{\text{release}}) = \tau'_r(F^0_{\text{ext}} - T)/\tau'\zeta$ is related to a second relaxation time $\tau'$.

Taken together these allow us to uniquely infer the parameters of (S.2a) from the the measured relaxation

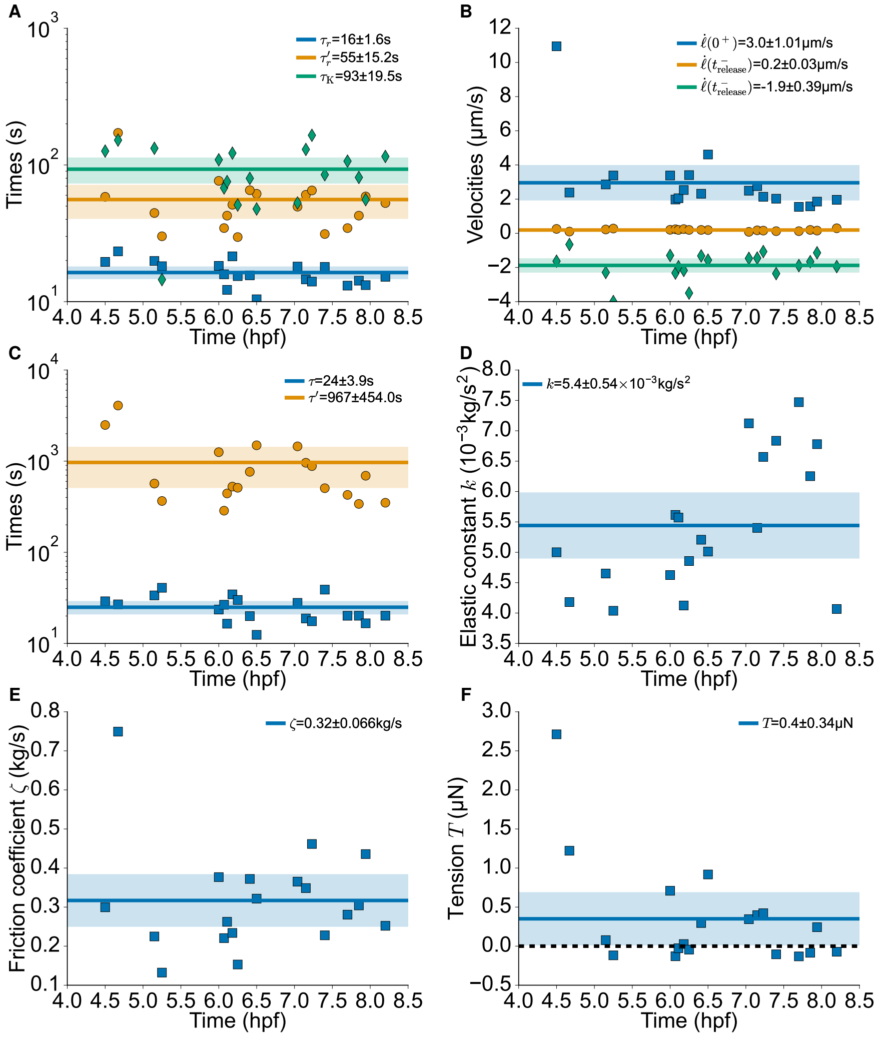


Figure S2: (A) Measured time scales $\tau_r$ and $\tau'_r$ for the relaxation of the velocities $\dot{\ell}(t)$ (Figure S1A) in the aspiration and release phases respectively, and measured time scale $\tau_K$ for the relaxation of the keratin $k(t)$ (Figure S1B). (B) Measured velocities as extracted by linear fits to the height curves $\ell(t)$ (Figure S1A) where $t_{\text{release}}$ is the time where the pressure is removed. (C) Computed relaxation time scales $\tau$ eq. (S.5a) and $\tau'$ (S.5b). (D) Tissue elastic constant $k$ eq. (S.5c). (E) Substrate friction coefficient $\zeta$ eq. (S.5d). (F) Internal tissue tension $T$ eq. (S.5e).

time scales and velocities (Figure S2A-B):

$$\tau = \frac{\tau_r \tau_r'}{\tau_r' - \tau_r \left(1 - \frac{\dot\ell(t^-_{\text{release}})}{\dot\ell(0^+)}\right)}, \tag{S.5a}$$

$$\tau' = \frac{\dot\ell(0^+)}{\dot\ell(t^-_{\text{release}})} \tau_r', \tag{S.5b}$$

$$k = \frac{F^0_{\text{ext}}}{\tau_r'(\dot\ell(t^-_{\text{release}}) - \dot\ell(t^+_{\text{release}}))} \left(1 - \frac{\dot\ell(t^-_{\text{release}})}{\dot\ell(0^+)}\right), \tag{S.5c}$$

$$\zeta = \frac{F^0_{\text{ext}}}{\dot\ell(t^-_{\text{release}}) - \dot\ell(t^+_{\text{release}})}, \tag{S.5d}$$

$$T = F^0_{\text{ext}} \left(\frac{\dot\ell(0^+)}{\dot\ell(t^-_{\text{release}}) - \dot\ell(t^+_{\text{release}})} - 1\right). \tag{S.5e}$$

We plot these quantities in Figure S2C-F.

##### Relation to the viscous drop model

Within the viscous drop model of Refs. [64, 65], the mechanical properties of the tissues are characterised in terms of surface tension $\gamma$, which provides the tension which the aspiration force must overcome, and viscosity $\eta$, which accounts for viscous dissipation at the opening of the pipette. Using our notations, these can be computed as follows [65]

$$\gamma = \frac{d\,\Delta P_{\text{ext}} \dot L_{\text{ret}}}{4(\dot L_{\text{ret}} + \dot L_{\text{asp}})} = \frac{T}{\pi d}, \tag{S.6a}$$

$$\eta = \frac{d\,\Delta P_{\text{ext}}}{6\pi(\dot L_{\text{asp}} + \dot L_{\text{ret}})} = \frac{2(\zeta + k\tau)}{3\pi^2 d}, \tag{S.6b}$$

where we have used the aspiration velocity $\dot L_{\text{asp}} = \dot\ell(t^-_{\text{release}}) = (F^0_{\text{ext}} - T)/(\zeta + k\tau)$ and the retraction velocity $\dot L_{\text{ret}} = -\dot\ell(t \gg t_{\text{release}}) = T/(\zeta + k\tau)$.

##### Keratin time scale

We expect the keratin concentration to respond to the stress within the cell. At the simplest level, *i.e.* without postulating a mechanism, we can extract a characteristic time scale for this response from the experimental curves, which we obtain by fitting their second part (after release) to an exponentially decreasing function $t \mapsto \exp(-t/\tau_K)$. This time scale $\tau_K$ is larger than the relaxation time scales $\tau_r$ and $\tau_r'$ (Figure S2A), consistent with the wider curves in the keratin than in the height plot (Figure S1).

#### 2 Vertex model and feedback

The results of the pipette experiment show that keratin reacts to stress, and that it also affects the elastic properties of the tissue. Together with the fact that keratin forms a growing network of filaments within cells during epiboly, we propose the following feedback mechanisms for keratin filament concentration:

(i) its concentration in a given cell increases with the inner pressure in this cell (mechanical) due to stress-dependent assembly and disassembly rates,

(ii) in turn the presence of keratin filaments generates a composite material, affecting the viscoelastic response of the cell by increasing both its stiffness (elasticity) and its relaxation time to external stress (viscosity); this effect is only considered above a minimum keratin concentration which translates the fact that keratin must percolate within the cell,

and implement these mechanisms within a vertex model.

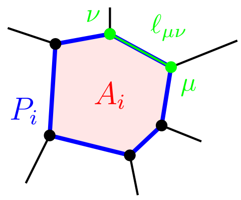


Figure S3: A cell in a vertex model.

#### Model formulation

We introduce a vertex model interaction potential with perimeter and area elasticity [68, 69]

$$U = \sum_{\text{cells } i} \left[ \frac{1}{2} \frac{\Gamma_i}{A_0} \left( A_i - A_i^0 \right)^2 + \frac{1}{2} \Gamma_i \left( P_i - s_0 \sqrt{A_i^0} \right)^2 \right], \tag{S.7}$$

where $A_i$ and $P_i$ are the area and perimeter of cell $i$ respectively (Figure S3), $\Gamma_i$ is an elastic constant, and $s_0 = P_i^0 / \sqrt{A_i^0}$ is the shape index. We introduce viscous relaxation through the cell-wise relaxation of the target area $A_i^0$

$$\tau_i \dot{A}_i^0 = -(A_i^0 - A_i) \tag{S.8}$$

while enforcing $A_i^0 = \max(A_0, A_i^0)$ with a reference area $A_0 > 0$ in order for cells not to disappear.

Mechanism (i), stress-dependent assembly of keratin filaments, is captured generically by the rate equation

$$\tau_K \dot{K}_i = \alpha \max(0, p_i) - K_i, \tag{S.9}$$

where $K_i$ and $p_i$ are the keratin concentration and the internal pressure (S.12) in cell $i$, and $\alpha > 0$ is the sensitivity of keratin to internal pressure. Under constant pressure ($p_i > 0$), the steady state keratin concentration is

$$K_i^{\text{ss}} = \alpha p_i \tag{S.10}$$

which is an increasing function of the pressure $p_i$. We consider initial keratin levels $K_i = 0$ for all cells.

The two-dimensional pressure of cell $i$ which derives from (S.7) is

$$p_i^{\text{2D}} = \frac{1}{A_i} \left[ \frac{\Gamma_i}{A_0} \left( A_i - A_i^0 \right) A_i + \Gamma_i \left( P_i - s_0 \sqrt{A_i^0} \right) P_i \right]. \tag{S.11}$$

To be consistent with the three-dimensional nature of the experiment, we introduce a modified three-dimensional pressure. Since cells spread enormously during epiboly, we introduce the volume $V_i = A_i h_i$ of the cell which we assume to be constant in time as in experiment, and which we set to an identical value for all cells, $V_i = V_0$. Then we write

$$p_i = p_i^{\text{3D}} = \frac{p_i^{\text{2D}}}{h_i} = \frac{1}{V_0} \frac{\Gamma_i}{A_0} (A_i - A_i^0) A_i, \tag{S.12}$$

where we have deliberately omitted the perimeter part of this quantity since the viscous relaxation (S.8) only depends on the area (other choices lead to unwelcome secondary feedback loops in the vertex model).

Mechanism (ii), the mechanical effect of keratin, is captured by adjusting stiffness and relaxation according to

$$\Gamma_i = \Gamma \ (1 + \beta \max(0, K_i - K_{\text{th}})), \tag{S.13a}$$

$$\tau_i = \tau \ (1 + \beta \max(0, K_i - K_{\text{th}})), \tag{S.13b}$$

where $\Gamma$ and $\tau$ are the keratin-free elastic constant and relaxation time scale respectively, $K_{\text{th}}$ is the threshold in keratin concentration, and $\beta$ characterises the intensity of the keratin effect on the cell properties. A positive threshold $K_{\text{th}} > 0$ is meant to translate the fact that keratin first needs to percolate through the cell for it to have an effect.

| Maxwell model | Vertex model |
|---|---|
| $\tau = 24\,\mathrm{s}$<br>$\tau' = 967\,\mathrm{s}$ | $\tau = 500\,\mathrm{s}$ |
| $k = 5.4 \times 10^3\,\mathrm{kg\,s^{-2}}$ | $\Gamma = 5.4 \times 10^3\,\mathrm{kg\,s^{-2}}$ |
| $\zeta = 0.32\,\mathrm{kg\,s^{-1}}$ | $\zeta = 0.32\,\mathrm{kg\,s^{-1}}$ |
| $T = 0.4\,\mu\mathrm{N}$<br>$F_{\mathrm{ext}} = 0.57\,\mu\mathrm{N}$ | $F_{\mathrm{YSL}} = 0.57\,\mu\mathrm{N}$ |
| | $s_0 = 3.72$ |
| | $A_0 = 600\,\mu\mathrm{m}^2$ |
| | $V_0 = 18 \times 10^3\,\mu\mathrm{m}^3$ |
| | $\alpha \sim \frac{K^{\mathrm{ss}}}{F_{\mathrm{YSL}/A_0}} \approx 2.4 \times 10^5\,\mathrm{kg^{-1}\,s^2}$ |
| | $\beta = 0$ (keratin loss-of-function), $\beta = 0.005$ (wild type) |

Table S1: Parameter values of the Maxwell model (Figure S2) and the vertex model.

#### Matching parameters and epiboly simulation

We finally write the equation of motion for vertex $\mu$

$$\zeta \dot{\boldsymbol{r}}_\mu = -\frac{\partial}{\partial \boldsymbol{r}_\mu} U + \boldsymbol{F}_\mu^{\mathrm{pull}}, \tag{S.14}$$

where $\boldsymbol{F}_\mu^{\mathrm{pull}}$ is an external pulling force that expands the tissue, simulating the effect of the YSL. We define boundary vertices as the ensemble of vertices at the open outer edge of the tissue. In order to keep the tissue round, we use the following pulling force

$$\boldsymbol{F}_\mu^{\mathrm{pull}} = \begin{cases} \frac{N_{\mathrm{edges}} F_{\mathrm{YSL}}}{\sum_\mu |\boldsymbol{\ell}_{\mu-1\to\mu} + \boldsymbol{\ell}_{\mu\to\mu+1}|} \left(\boldsymbol{\ell}_{\mu-1\to\mu} + \boldsymbol{\ell}_{\mu\to\mu+1}\right) \times \hat{\boldsymbol{e}}_z & \text{if } \mu \text{ is a boundary vertex,} \\ 0 & \text{otherwise,} \end{cases} \tag{S.15}$$

where $N_{\mathrm{edges}}$ is the number of boundary vertices, indexed by $\mu$ in anticlockwise order, with $\boldsymbol{\ell}_{\mu\to\nu} = \boldsymbol{r}_\nu - \boldsymbol{r}_\mu$.

We estimate the model parameters (Table S1) as follows: We use the same elastic constant $k = \Gamma$ and friction coefficient $\zeta$. We use a relaxation time scale $\tau$ for the vertex model with a value intermediate between the two time scales $\tau$ and $\tau'$ of the Maxwell model. We fix the shape index $s_0 = 3.72$ so that the tissue is in a solid state in the absence of viscous relaxation [69]. We estimate the reference cell area to $A_0 = 600\,\mu\mathrm{m}^2$ and the reference cell height $h_0 = 30\,\mu\mathrm{m}$ so that the reference cell volume $V_0 = h_0 A_0 = 18 \times 10^3\,\mu\mathrm{m}^3$.

The action of the YSL is simulated through a force ramp from 0 to a value of $F_{\mathrm{YSL}} = F_{\mathrm{ext}}$, consistent with values for the internal tissue tension $T$ in the pipette model that average just below this value. We estimate the coefficient $\alpha$ by considering a reference value $K^{\mathrm{ss}} \approx 225$ (main text Figure 1(c)) and a reference pressure $p \sim F_{\mathrm{YSL}}/A_0$, such that using (S.10) we obstain $\alpha \sim K^{\mathrm{ss}} A_0 / F_{\mathrm{YSL}} \approx 2.4 \times 10^5\,\mathrm{kg^{-1}\,s^2}$. Note that the keratin concentrations in the pipette experiment cannot be compared to the epiboly experiments directly due to differences in imaging. Then $\beta$ remains as a free parameter which we can adjust to match the experimental observations.

We generate initial disordered configurations by applying an active Brownian force $\boldsymbol{F}_\mu^{\mathrm{AB}}$ on all vertices and an additional boundary tension force $\boldsymbol{F}_\mu^{\mathrm{BT}}$ on boundary vertices to keep the tissue round. We define the active Brownian force

$$\boldsymbol{F}_\mu^{\mathrm{AB}} = v_0 \begin{pmatrix} \cos\theta_\mu \\ \sin\theta_\mu \end{pmatrix}, \tag{S.16a}$$

$$\dot{\theta}_\mu = \sqrt{1/\tau_p} \eta_\mu, \tag{S.16b}$$

where $\eta_\mu$ a Gaussian white noise with variance $\langle \eta_\mu(0)\eta_\nu(t)\rangle = \delta_{\mu\nu}\,\delta(t)$, and the boundary tension force

$$\boldsymbol{F}_\mu^{\rm BT} = \gamma\left(\frac{\boldsymbol{\ell}_{\mu\to\mu-1}}{|\boldsymbol{\ell}_{\mu\to\mu-1}|} + \frac{\boldsymbol{\ell}_{\mu\to\mu+1}}{|\boldsymbol{\ell}_{\mu\to\mu+1}|}\right). \tag{S.17}$$

which only acts on outer boundary vertices. We used the interaction potential (S.7) with parameters $\Gamma_i = 1$, $A_i^0 = 1$, $s_0 = 3.72$, the active Brownian force (S.16) with parameters $v_0 = 0.1$, $\tau_p = 1$, and the boundary tension force (S.17) with parameter $\gamma = 0.3$. We perform a first initial run to reach steady state, then remove the active Brownian force and perform a second run to reach a force equilibrium between the interaction forces (S.7) and the boundary tension forces (S.17). We use this final configuration to initialise the simulation with the parameters from Table S1. We rescale the size of the system to minimise $|\sum_{\text{vertices }\mu}(\partial_{\boldsymbol{r}_\mu}U)\cdot(\boldsymbol{r}_\mu - \boldsymbol{r}^{\rm CM})|$, where $\boldsymbol{r}^{\rm CM} = \sum_{\text{vertices }\mu}\boldsymbol{r}_\mu/N_{\rm v}$ is the position of the centre of mass and $N_v$ the number of vertices. This latter rescaling ensures that the tissue does not initially move unless it is being pulled.

### 3 Model tissue cell ablation

We prepare model tissue cell ablation simulations by first generating a disordered configuration of the vertex model with periodic boundary conditions. To this effect we use the potential energy (S.7) with parameters $\Gamma_i = 1$, $A_i^0 = 1$, $s_0 = 3.81$ and the active Brownian force (S.16) with parameters $v_0 = 0.25$, $\tau_p = 1$. We perform a first initial run to reach steady state. We then remove the active Brownian force and rescale the system such that the mean area of the cells is $\overline{A} = 1750\,\mu\text{m}^2$, and set the other parameters to the values in Table S1. We set the target areas of the cells at $A_i^0 = A_i/2.02$ but keep $\tau_i = \infty$ in (S.8), and let the system equilibrate to a steady state in keratin intensity. At $t = 0$ we set $\tau_i = 500\,\text{s}$, ablate 6 cells, and apply a boundary force (S.17) around the ablated cells with $\gamma = 3\,\mu\text{N}$.

### 4 Keratin feedback regulates velocity

To understand the keratin expression dynamics and edge speeds, we turn to simple mean field dynamical equations. If we return to the dynamics of a single junction, now as part of epiboly, we have

$$\zeta\dot{\ell} = -k(\ell - \ell_0) + F_{\rm YSL}, \quad \tau\dot{\ell}_0 = \ell - \ell_0, \tag{S.18}$$

with keratin dynamics

$$\tau_K\dot{K} = \alpha\max(0, p) - K. \tag{S.19}$$

The keratin then influences the stiffness and relaxation constants equally through

$$k = k_0(1 + \beta\max(0, K - K_{th})), \quad \tau = \tau_0(1 + \beta\max(0, K - K_{th})). \tag{S.20}$$

Here, to match the $p^{\rm 3D}$ in the vertex model, we write the tissue pressure $p = k\frac{(\ell-\ell_0)\ell}{V_0}$ and we subsequently use the same parameters as in the vertex model.

Figure S4 shows the dynamics of this mean field model. As in the main text, we perform a linear force ramp over 11 hours for different values of $F_{\rm YSL}$, both above and below the expected value of $0.57\mu N$ for a wild-type embryo. Panel B show the keratin dynamics, which is quantitatively close to the full model and the experiment. Panel C shows the rate of junction expansion $\dot{\ell}$. As in the full model, it quickly rises proportional to pulling force, and then has a peak when the feedback kicks in when $K \geq K_{\rm th}$. Junction expansion rate $\dot{\ell}$ can be related to the edge speed through the scaling $v_{\rm edge} \approx \sqrt{N}\dot{\ell}$, i.e. multiply by a factor of 23. Again, the dynamics is quantitatively similar to the full vertex model, though the peak is sharper. The smoothing is therefore due to the strong heterogeneity in keratin expression in the full simulation. If we turn off the feedback ($\beta = 0$), just as in the vertex model, the rate of junction expansion is larger and increases indefinitely instead (panel D). In summary, the mean field model captures the essential features of the full vertex model.

We can understand the origin of the feedback regulation in this simplified setting: In epiboly, we expect pressure to be positive, i.e. a tissue under tension, so we can replace $\max(0, p) = p$. We can then

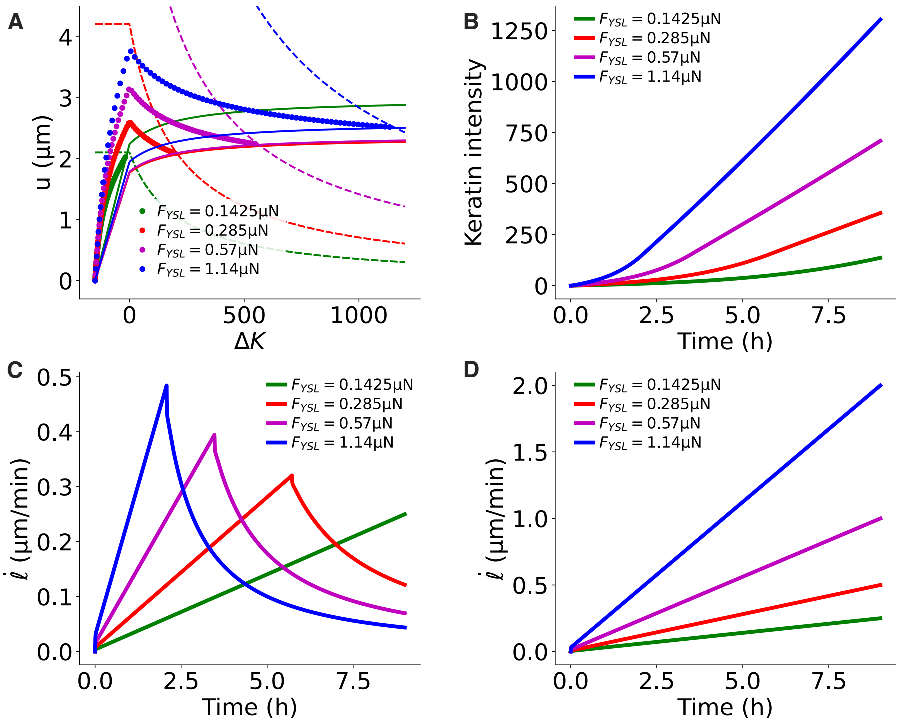


Figure S4: Mean-field model dynamics. As in the main text, we perform a linear force ramp for four values of $F_{\rm YSL}$. (A) Reduced variable phase space $(\Delta K, u)$. Simulations with feedback ($\beta = 0.005$) are shown as dots, together with the nullclines of $\Delta K$ (solid) and of $u = \ell - \ell_0$ (dashed). (B) Keratin dynamics and (C) junction length expansion speed $\dot{\ell}$ as a function of time. (D) For the 'mutant' model with no feedback ($\beta = 0$), $\dot{\ell}$ is not regulated and increases with $F_{\rm YSL}$.

define the new variables $\Delta K = K - K_{th}$ and $u = \ell - \ell_0$, the current deformation from the mechanical equilibrium value. For $\Delta K > 0$ we then write

$$\dot{u} = -\left[\frac{k_0}{\zeta}(1+\beta\Delta K) + \frac{1}{\tau_0(1+\beta\Delta K)}\right]u + \frac{F_{\rm YSL}}{\zeta} \tag{S.21a}$$

$$\Delta\dot{K} = \frac{\alpha k_0 a}{V_0}(1+\beta\Delta K)u - \frac{1}{\tau_K}(\Delta K + K_{th}) \tag{S.21b}$$

and the same equation with $\beta = 0$ for $\Delta K < 0$. We have used $\ell(\ell - \ell_0) \approx a\,u$, where $a = \mathcal{O}(\ell)$ is a constant length scale in the model, under the assumption that $\ell$ is slowly varying. This slight simplification makes the model tractable.

We are mostly interested in the long-term steady-state evolution set by the fixed points of the dynamics, where $\dot{K} = 0$ and $\dot{u} = 0$ and so the tissue expands at constant keratin concentration. The fixed point (and there is only one) is set by the intersections of the nullclines, i.e. the lines where separately $\dot{u} = 0$ (u-nullcline) and $\dot{K} = 0$ (K-nullcline). Their equations are given by:

$$u_{\dot{u}=0} = \frac{F_{\rm YSL}\tau_0}{\zeta}\frac{1+\beta\Delta K}{1+\frac{k_0\tau_0}{\zeta}(1+\beta\Delta K)^2} \tag{S.22a}$$

$$u_{\dot{K}=0} = \frac{\Delta K + K_{th}}{\frac{\alpha k_0 a}{V_0}(1+\beta\Delta K)} \tag{S.22b}$$

for $\Delta K > 0$ and the same with $\beta = 0$ for $\Delta K < 0$. The edge speed is given by $\dot{\ell} = \dot{u} + \dot{\ell}_0 = -\frac{u^*}{\tau(\Delta K^*)}$, where the last equality is valid at the fixed point $(K^*, u^*)$.

Figure S4A shows the simulated dynamics replotted in $(\Delta K, u)$ coordinates. We have added the nullclines of the keratin dynamics $\Delta K$ (solid lines) and of the junction dynamics $u$ (dashed lines), both plotted for the end values of the simulations for $F_{\mathrm{YSL}}$ and $a$. Here the junction nullcline moves up and left with increasing pulling force. The keratin nullcline depends only slightly on $\ell$, in contrast.

The following picture emerges: At low pulling forces, i.e. either early in the simulation or if $F_{\mathrm{YSL}}$ is small, the junction nullcline intersects the keratin nullcline for $\Delta K < 0$, where it is linearly increasing as a function of $\Delta K$. This corresponds to the combined upwards ramp of keratin and junction speed that we observe at early times, and to a model with no feedback. At higher force, the junction nullcline intersects the keratin nullcline at $\Delta K > 0$, where the curve becomes nearly flat. Thus we find the characteristic dynamics of feedback: Independent of pulling force, we recover approximately the same $u^*$ at the fixed point, while $\Delta K^*$ keeps increasing with pulling force. This translates to reducing $\dot{\ell}$ with pulling force as $\dot{\ell} = -\frac{u^*}{\tau(\Delta K^*)}$ and $\tau$ increases with $\Delta K^*$.

In contrast, when we turn off feedback ($\beta = 0$), $\dot{\ell}$ keeps increasing indefinitely with pulling force, and one can easily compute that $\dot{\ell} = \frac{F_{\mathrm{YSL}}}{\zeta + k_0 \tau_0}$ at the fixed point.
